# TMEM165 replenishes lysosomal Ca^2+^ stores, protects cells against Ca^2+^ overload, and mediates Ca^2+^-induced lysosomal H^+^ leakage

**DOI:** 10.1101/2024.05.09.593345

**Authors:** Ran Chen, Bin Liu, Dawid Jaślan, Lucija Kucej, Veronika Kudrina, Belinda Warnke, Yvonne Klingl, Arnas Petrauskas, Kenji Maeda, Christian Grimm, Marja Jäättelä

## Abstract

The proper function of lysosomes depends on their ability to store and release calcium. While several lysosomal calcium release channels have been described, how mammalian lysosomes replenish their calcium stores has not been determined. Using genetic depletion and overexpression techniques combined with electrophysiology and visualization of subcellular ion concentrations and their fluxes across the lysosomal membrane, we show here that TMEM165 imports calcium to the lysosomal lumen and mediates calcium-induced lysosomal proton leakage. Accordingly, TMEM165 accelerates the recovery of cells from cytosolic calcium overload thereby enhancing cell survival while causing a significant acidification of the perilysosomal area and the entire cytosol. These data indicate that in addition to its essential role in the glycosylation in the Golgi, a small but significant fraction of TMEM165 localizes on the lysosomal limiting membrane, where it protects cells against cytosolic calcium overload, preserves lysosomal function by refilling lysosomal calcium stores and regulates perilysosomal and lysosomal pH.

## Introduction

Calcium (Ca^2+^) and proton (H^+^) homeostasis are of crucial importance for virtually all aspects of cellular life. In mammalian cells, the free Ca^2+^ concentration in the cytosol [Ca^2+^]_cyt_ is maintained approximately four orders of magnitude lower than that in the extracellular space and the major intracellular Ca^2+^ storage organelles ^1, 2^. Similarly, the concentration of protons in the cytosol [H^+^]_cyt_ is up to three orders of magnitude lower than that in the lumen of lysosomes ^2^. Such steep ion concentration gradients across cellular membranes are maintained by numerous pumps, and exchangers that remove excess Ca^2+^ and H^+^ from the cytosol to their storage organelles or the extracellular space ^1, 3, 4^. Due to the effective cytosolic buffering systems for both ions, cells tolerate transient fluctuations in their concentrations. In fact, such fluctuations are essential for a wide range of physiological signaling cascades ^1, 5^. Greater and more sustained changes in ion homeostasis are, however, associated with pathological processes, and prolonged increases in either [Ca^2+^]_cyt_ or [H^+^]_cyt_ eventually lead to cell death and organ dysfunction ^6, 7^.

Late endosomes/lysosomes (hereafter referred to as lysosomes) are membrane-surrounded intracellular organelles originally described as sites of degradation and recycling of cellular organelles and macromolecules ^8, 9^. In recent years, lysosomes have emerged as versatile signaling organelles that regulate numerous cellular functions from metabolic pathways and cell division to malignant transformation and cell death ^10–13^. Most, if not all, lysosomal activities depend on ion channels and transporters that establish electrolyte concentration gradients and membrane potential across the lysosomal limiting membrane ^2, 14^. Due to the low pH optimum of most lysosomal hydrolases, the catabolic function of lysosomes depends on the vacuolar-type H^+^-ATPase (V-ATPase), which continuously pumps protons from the cytosol to the lysosomal lumen ^15, 16^. On the other hand, TMEM175, a H^+^-activated and H^+^-permeable channel in the lysosomal limiting membrane, protects lysosomes from overacidification by leaking H^+^ in response to low luminal pH ^17, 18^. The significant alkalization of TMEM175-depleted lysosomes observed upon V-ATPase inhibition suggests that TMEM175 is not the only protein capable of releasing H^+^ from the lysosomal lumen ^17^.

The endoplasmic reticulum (ER), Golgi, and lysosomes constitute major intracellular Ca^2+^ stores that can release Ca^2+^ to the cytosol in response to various second messengers activated by intra- and extracellular cues ^1, 19, 20^. While the sarco/endoplasmic reticulum Ca^2+^ ATPases (SERCAs) constantly pump free cytosolic Ca^2+^ to the ER, and SERCA and secretory pathway Ca^2+^ ATPases pump it to the Golgi, the mechanisms by which mammalian lysosomes replenish their Ca^2+^ stores remain largely unknown. In plant and yeast cells, the Ca^2+^ filling of their lysosome-related organelles, vacuoles, is mediated by Ca^2+^/H^+^ exchangers CAX and VCX1, respectively ^21, 22^. Although strong experimental evidence suggests that a similar Ca^2+^/H^+^ exchanger activity drives lysosomal Ca^2+^ filling in mammalian cells, CAX and VCX1 have neither mammalian orthologs nor known functional counterparts ^20, 21^. Given that the V-ATPase constitutively pumps cytosolic H^+^ into the lysosomal lumen, the resulting steep H^+^ gradient across the lysosomal membrane could in theory provide the driving force for a putative lysosomal Ca^2+^/H^+^ exchanger. Such an exchanger could also explain the TMEM175-independent lysosomal H^+^ leakage discussed above and the cytosolic acidification observed in response to elevated [Ca^2+^]_cyt_ observed for example in glutamate-treated rat sensory neurons ^23^, ionomycin-treated rabbit corneal epithelial cells ^24^, rat parotid and pancreatic acinar cells treated with various Ca^2+^ mobilizing agents ^25, 26^, and human cancer cells treated with cationic amphiphilic drugs (CADs) that trigger lysosomal Ca^2+^ release via the purinergic receptor P2X4 (P2RX4) and subsequent lysosome-dependent cancer cell death ^27^.

In living cells, the lysosomal H^+^ leakage has hitherto been demonstrated only indirectly by measuring changes in the lysosomal and cytosolic pH. To detect the lysosomal H^+^ leakage directly, we created a spatially restricted pH sensor that allowed us to measure local pH changes in the perilysosomal area. Using this tool, we confirmed the TMEM175-mediated basal lysosomal H^+^ leakage in living cells and demonstrated that TMEM175-independent perilysosomal acidification occurs in response to various Ca^2+^ mobilizing agents. The latter triggered a search for a lysosomal Ca^2+^/H^+^ exchanger and the identification of TMEM165, which has been previously described as a putative cation/H^+^ antiporter that exchanges Ca^2+^ and manganese (Mn^2+^) for H^+^ across the Golgi membrane ^28^, as a mediator of Ca^2+^-induced lysosomal H^+^ leakage in mammalian cells. Taken together, our data provide a molecular explanation for the long-sought Ca^2+^ influx pathway into lysosomes.

## Results

### Lysosomal proton leakage acidifies the perilysosomal area

To visualize lysosomal H^+^ efflux in living cells, we constructed a cDNA encoding for a perilysosomal pH sensor consisting of 39 amino-terminal amino acids of late endosomal/lysosomal adaptor, MAPK and MTOR activator 1 (LAMTOR1), which target the protein to the lysosomal surface ^29^, and a pH-sensitive fluorescent protein, mKeima, whose fluorescence intensity negatively correlates with the adjacent pH ^30^ (Fig. 1a,b). In stably transfected HeLa human cervix carcinoma cells, the obtained Lyso-mKeima sensor colocalized mainly with established lysosomal markers, Lysotracker® green (LTG) and monomeric green fluorescence (mGFP)-tagged lysosomal associated membrane protein 1 (LAMP1) but not with the Golgi apparatus visualized by enhanced GFP (EGFP)-tagged RAB6A ^31^, ER visualized by ER-tracker^TM^ Green or mitochondria visualized by Mitotracker^TM^ Green (Fig. 1a and Extended Data Fig. 1a-c). Using Lyso-mKeima and either SypHer3s or pHrodo^TM^-Green-AM to detect perilysosomal and cytosolic pH, respectively, we demonstrated that the H^+^ concentration on the lysosomal surface of HeLa cells (pH 7,05 ± 0.01) was 2.9-fold higher than that in the cytosol (pH 7.52 ± 0.03) (Fig. 1c). Substantial acidification of the perilysosomal area was also observed in all the other human cell lines tested, *i.e.* in MCF7 (2.2-fold), MDA-MB-231 (1.6-fold) and MDA-MB-468 (2.1-fold) breast carcinoma cells and in noncancerous MCF10A breast epithelial cells (1.6-fold) (Fig. 1c and Extended Data Fig. 1d). In further support of the lysosomal H^+^ leakage as the cause of the perilysosomal acidification, siRNA-mediated partial depletion of the lysosomal H^+^-activated H^+^ channel TMEM175 in HeLa-Lyso-mKeima cells significantly increased the perilysosomal pH (Fig. 1d,e). Further supporting the existence of TMEM175-mediated basal lysosomal H^+^ leakage, Lyso-mKeima fluorescence increased slightly but significantly upon concanamycin A-mediated inhibition of the V-ATPase in control cells but not in TMEM175-depleted cells (Fig. 1f). Even stronger increases in Lyso-mKeima fluorescence intensities were observed upon treatment of HeLa cells with arachidonic acid (Fig. 1g), which has been shown to induce a TMEM175-dependent decrease in cytosolic pH (pH_cyt_) ^17^. In contrast to the reported complete inhibition in TMEM175 knockout (KO) cells ^17^, TMEM175 siRNAs only partially inhibited the lysosomal H^+^ efflux induced by arachidonic acid (Fig. 1h).

**Figure 1.**
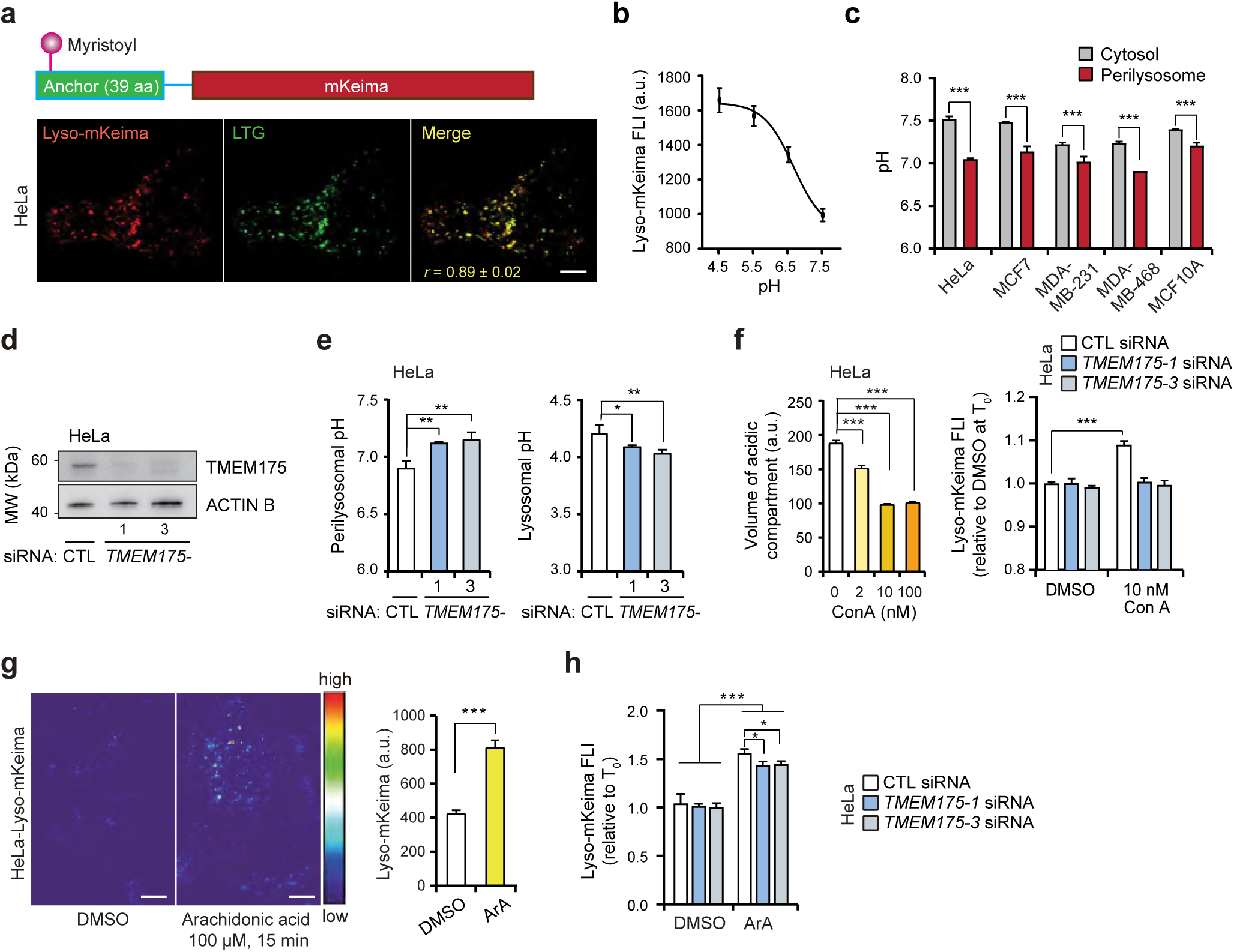
TMEM175 acidifies the perilysosomal area. a. Construct of Lyso-mKeima (top). Representative (n =3) confocal images of live HeLa-Lyso-mKeima stained with Lysotracker® Green (bottom). r, Pearson’s colocalization coefficient (n =20). Scale bars, 10 µm. b. The standard curve for fluorescence intensity (FLI) of mKeima at pH ranging from 4.5 to 7.5. c. Cytosolic and perilysosomal pH values of indicated cell lines. d. Representative (n=3) immunoblots of indicated proteins in HeLa cells treated with indicated siRNAs for 72 h. e. Perilysosomal pH analyzed by Lyso-mKeima (left) and lysosomal pH analyzed by LysoSensorTM Yellow/Blue FLI (right) in HeLa cells treated with indicated siRNAs for 72 h. f. Relative volumes of acidic compartments in HeLa cells treated with Con A for 1 h, stained with Lysotracker green, and analyzed by flow cytometry (left). mKeima FLI in HeLa-Lyso-mKeima cells treated for 72 h with indicated siRNAs and with either DMSO or 10 nM Con A for 1 h (right). g. Representative (n = 3) images of live HeLa-Lyso-mKeima cells treated with 100 µM Arachidonic acid for 15 min (left). mKeima FLI of cells shown on left (right). h. mKeima FLI in HeLa-Lyso-mKeima cells treated for 72 h with indicated siRNAs and with either DMSO or 100 µM Arachidonic acid for 20 min. Error bars, SD of three independent experiments with ≥10 (a bottom, b, c, e, f right, g, h), or ≥10000 (f left) randomly chosen cells analyzed in each sample. *, P < 0.05; **, P < 0.01; ***, P < 0.001 as analyzed by one-way Anova (e, f left, g right) or two-way Anova (c, f right, h) with Tukey (e, f left, g right) or Dunnett (c, f right, h) multiple comparison.

Taken together, these data introduce Lyso-mKeima as a useful tool to detect perilysosomal pH and lysosomal H^+^ leakage and demonstrate that TMEM175-mediated basal lysosomal H^+^ leakage maintains the pH of the perilysosomal area significantly below the value of the average cytosolic pH.

### An increase in [Ca^2+^]_cyt_ triggers lysosomal H^+^ leakage

Commonly used cationic amphiphilic drugs (CADs) including antihistamines, antidepressants and antipsychotics are emerging as potent anticancer drugs ^12^. As basic and membrane permeable molecules, they accumulate in the lysosomal lumen, where they inhibit lysosomal function and induce lysosomal membrane permeabilization specifically in cancer cells ^32, 33^. We have previously shown that treating cancer cells with CADs induces cytosolic acidification hours before any signs of lysosomal membrane damage or cell death occur ^34^. Supporting lysosomal H^+^ efflux as the mechanism of CAD-induced cytosolic acidification, the CAD antihistamines, ebastine and terfenadine, triggered rapid and significant increases in the Lyso-mKeima fluorescence intensity (Fig. 2a,b). Importantly, these increases originated from the entire cell population and from the presence of dozens of Lyso-mKeima positive puncta *per* cell rather than from a few dying cells or individual damaged lysosomes (Fig. 2a). Notably, CAD-induced lysosomal H^+^ flux was insensitive to depletion of TMEM175 (Fig. 2c).

**Figure 2.**
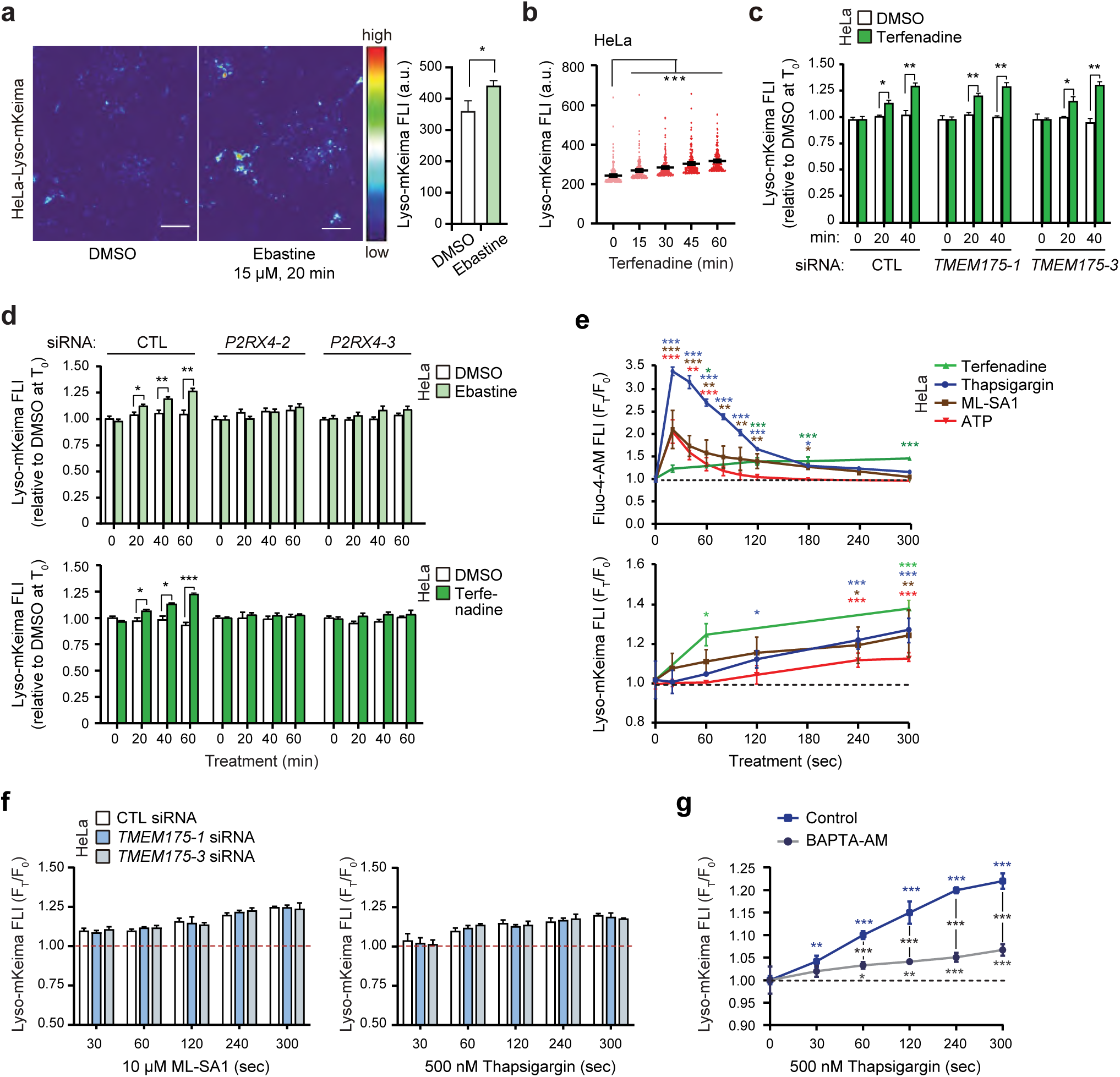
Ca2+ mobilizing agents induce TMEM175-independent lysosomal H+ leak. a. Representative (n = 3) images of live HeLa-Lyso-mKeima cells treated with 15 µM ebastine for 20 min (left). mKeima FLI of cells shown on left (right). b. Kinetics of mKeima FLI in HeLa cells treated with 6 µM terfenadine. c. mKeima FLI in HeLa-Lyso-mKeima cells treated for 72 h with indicated siRNAs and with either DMSO or 6 µM terfenadine for the last 20 or 40 min. d. mKeima FLI in HeLa-Lyso-mKeima cells treated for 72 h with indicated siRNAs and with 15 µM ebastine (top) or 6 µM terfenadine (bottom) for the last 20, 40 or 60 min. e. Fluo-4-AM FLI (top) and mKeima FLI (bottom) in HeLa cells treated with 6 µM terfenadine, 500 nM thapsigargin, 10 µM ML-SA1 or 1 µM ATP for the last 0 – 300 sec and analyzed by flow cytometry. f. mKeima FLI in HeLa-Lyso-mKeima cells treated for 72 h with indicated siRNAs and with 10 µM ML-SA1 (left) or 500 nM thapsigargin (right) for the last 0 – 300 sec. g. mKeima FLI in HeLa-Lyso-mKeima cells pretreated for 30 min with 20 µM BAPTA-AM and treated with 500 nM thapsigargin for the last 0 – 300 sec. Error bars, SD of three independent experiments with ≥10 (a, b, c, d, f, g), or ≥10000 (e) randomly chosen cells analyzed in each sample. *, P < 0.05; **, P < 0.01; ***, P < 0.001 as analyzed by one-way Anova (a, b) or two-way Anova (c, d, e, f, g) with Tukey (a, b) or Dunnett (c, d, e, f, g) multiple comparison.

To determine the regulatory mechanism of CAD-induced, TMEM175-independent lysosomal H^+^ leakage, we first examined the role of CAD-induced rapid Ca^2+^ release in this process. Interestingly, siRNA-mediated depletion of P2RX4, a Ca^2+^ channel responsible for the CAD-induced lysosomal Ca^2+^ release ^27^, effectively blocked ebastine- and terfenadine-induced perilysosomal acidification in HeLa-Lyso-mKeima cells (Fig. 2d and Extended Data Fig. 2a), while neither the slightly acidic pH on the surface of lysosomes in control siRNA-treated cells (Extended Data Fig. 2b) nor its further acidification by maximal inhibition of the V-ATPase depended on P2RX4 (Extended Data Fig. 2c).

The reliance of CAD-induced lysosomal H^+^ leakage on the P2RX4 Ca^2+^ channel inspired us to further explore the link between lysosomal Ca^2+^ release and perilysosomal acidification. Given that the elevation in the [Ca^2+^]_cyt_ controls diverse cellular functions and that lysosomal Ca^2+^ uptake has been reported to depend on acidic lysosomal pH ^21, 35^, we investigated the possibility that lysosomes export H^+^ in exchange for importing excessive cytosolic Ca^2+^. For this purpose, we treated the Lyso-mKeima-expressing HeLa cells with three stimuli that increase the [Ca^2+^]_cyt_ via different means: (i) thapsigargin, which inhibits the ER and Golgi resident SERCA Ca^2+^ pumps ^36^, (ii) ATP, which stimulates P2 purinoreceptors to generate inositol 1,4,5-triphosphate (IP_3_), which in turn triggers the release of Ca^2+^ from the ER through IP_3_ receptor-regulated channels ^37^, and (iii) ML-SA1, an agonist of the lysosomal Ca^2+^ release channel, mucolipin TRP cation channel 1 (TRPML1) ^38^.

Following rapid and transient increases in the [Ca^2+^]_cyt_ (Fig. 2e, *top*), all three treatments induced significant increases in Lyso-mKeima fluorescence intensity, which was indicative of decreased perilysosomal pH and lysosomal H^+^ leakage (Fig. 2e, *bottom*). In contrast to the transient [Ca^2+^]_cyt_ peak induced by thapsigargin, ATP, and ML-SA1, the Ca^2+^ peak induced by CADs was weaker and longer lasting (Fig. 2e, *top*) ^27^, and was followed by a more substantial and persistent decrease in perilysosomal pH (Fig. 2e, *bottom* and Extended Data Fig. 2d). Like CADs, thapsigargin and ML-SA1 induced TMEM175 independent lysosomal H^+^ effluxes (Fig. 2f). However, the lysosomal H^+^ efflux triggered by thapsigargin was effectively inhibited by pretreatment of cells with the intracellular Ca^2+^ chelator, BAPTA-AM (Fig. 2g).

The data presented above corroborate the ability of excess cytosolic Ca^2+^ to trigger lysosomal H^+^ leakage.

### TMEM165 mediates Ca^2+^-induced lysosomal proton leakage

In our search for a protein mediating Ca^2+^ influx into lysosomes, transmembrane protein 165 (TMEM165), a putative divalent cation/H^+^ exchanger with a well-described role in manganese homeostasis and glycosylation of proteins and lipids in the Golgi network ^28, 39–41^, attracted our attention. In addition to its presence in the Golgi apparatus, TMEM165 has been detected in the lysosomal compartment of HeLa cells, where its depletion has been reported to increase the volume of the acidic compartment ^28, 42^. Supporting the role of TMEM165 in CAD-induced, lysosomal H^+^ leakage, partial depletion of this gene by two independent siRNAs effectively reduced ebastine- and terfenadine-induced lysosomal H^+^ leakage and subsequent acidification of the cytosol in HeLa cells (Fig. 3a-c). The strong TMEM165 dependences of CAD-induced lysosomal H^+^ leakage and cytosolic acidification were also observed in MDA-MB-468 and MCF7 cells, respectively (Extended Data Fig. 3a-f). The depletion of *TMEM165* in HeLa and MDA-MB-468 cells also inhibited lysosomal H^+^ leakage induced by other Ca^2+^-mobilizing agents, ML-SA1 and thapsigargin (Fig. 3d and Extended Data Fig. 3g), while TMEM165 was dispensable for the perilysosomal acidification induced by arachidonic acid and concanamycin A (Extended Data Fig. 3h,i). Furthermore, siRNA-mediated depletion of TMEM165 had no effect on the perilysosomal pH in untreated HeLa cells (Fig. 3e), and even complete CRISPR-mediated depletion of TMEM165 in either HeLa or MDA-MB-468 cells altered neither the lysosomal nor the cytosolic pH in unstimulated cells (Fig. 3f-h and Extended Data Fig. 3e,f,j). However, the overexpression of either TMEM165-Flag or TMEM165-mCherry in HeLa cells significantly altered the intracellular pH homeostasis by increasing the lysosomal pH and acidifying not only the perilysosomal area but also the entire cytosol (Fig. 3e-i).

**Figure 3.**
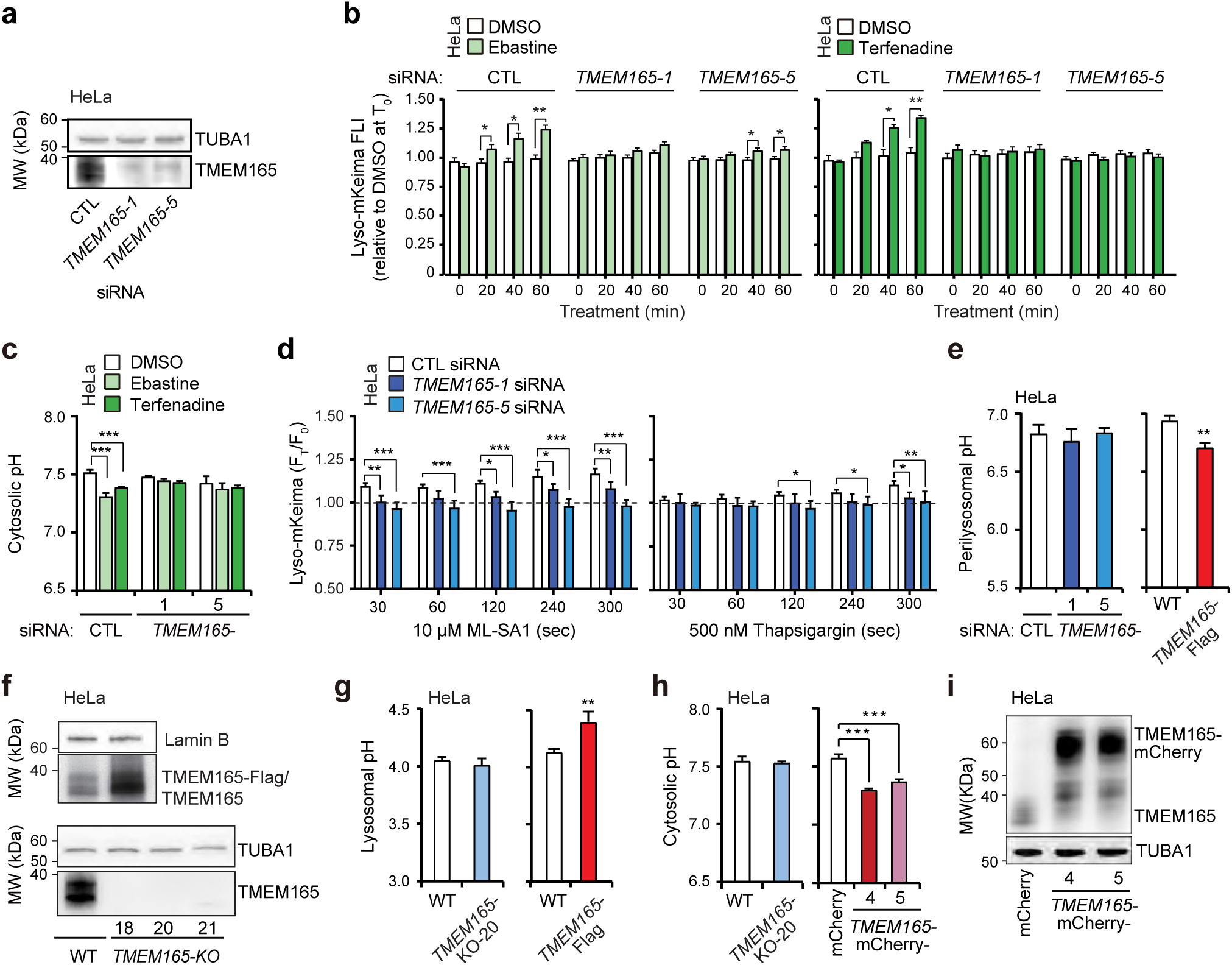
Ca2+-induced lysosomal H+ leak is mediated by TMEM165. a. Representative (n=3) immunoblots of indicated proteins in HeLa cells treated with indicated siRNAs for 72 h. b. mKeima FLI in HeLa-Lyso-mKeima cells treated for 72 h with indicated siRNAs and with 15 µM ebastine (left) or 6 µM terfenadine (right) for the last 20, 40 or 60 min. c. Cytosolic pH analyzed by pHrodo-AM FLI in HeLa cells treated with indicated siRNAs for 72 h and with 15 µM ebastine or 6 µM terfenadine for 2 h. d. mKeima FLI in HeLa-Lyso-mKeima cells treated for 72 h with indicated siRNAs and with 10 µM ML-SA1 (left) or 500 nM thapsigargin (right) for the last 0 – 300 sec. e. Perilysosomal pH analyzed by Lyso-mKeima FLI in HeLa cells treated with indicated siRNAs for 72 h (left) or in TMEM165 overexpressed cells (right) f. Representative (n=3) immunoblots of indicated proteins in WT, TMEM165 overexpressed cell lines (top) and TMEM165-KO cell lines (bottom). g. Lysosomal pH analyzed by LysoSensorTM Yellow/Blue FLI in HeLa TMEM165-KO cells (left) and TMEM165 overexpressed cells (right). h. Cytosolic pH analyzed by pHrodo-AM FLI in HeLa TMEM165-KO cells (left) and TMEM165 overexpressed cells (right). i. Representative (n=3) immunoblots of indicated proteins in mCherry- and TMEM165-mCherry-trasfected HeLa clones. Error bars, SD of three independent experiments with ≥10 randomly chosen cells analyzed in each sample. *, P < 0.05; **, P < 0.01; ***, P < 0.001 as analyzed by one-way Anova (e left, h right) or two-way Anova (b, c, d) with Tukey (e left, h right) or Dunnett (b, c, d) multiple comparison. (e right, g, h left) were analyzed by unpaired t-test.

These data indicate that TMEM165 is required for the lysosomal H^+^ leakage in response to elevations in the free [Ca^2+^]_cyt_, and that its overexpression disturbs cellular pH homeostasis.

### TMEM165 replenishes lysosomal Ca^2+^ stores and protects cells against Ca^2+^ mobilizing agents

Next, we investigated the effect of TMEM165 on the cellular Ca^2+^ homeostasis. To determine the effect of TMEM165 on the capacity of lysosomes to store Ca^2+^, we measured the fluorescence intensity (FLI) of a cytosolic Ca^2+^ indicator Fluo4-AM following the mobilization of lysosomal Ca^2+^ with glycyl-L-phenylalanine-beta-naphthylamide (GPN), a lysosomotrope that permeabilizes lysosomal membranes in a cathepsin C-dependent manner ^43, 44^. The increase in the free [Ca^2+^]_cyt_ upon lysosomal membrane permeabilization was significantly reduced or increased in TMEM165-depleted and TMEM165-Flag expressing HeLa cells, respectively (Fig. 4a,b). Similarly, GPN-mediated lysosomal membrane permeabilization significantly increased the free [Ca^2+^]_cyt_ less in MDA-MB-468-TMEM165-KO cells than in parental MDA-MB-486 cells (Fig. 4c). In further support of the ability of TMEM165 to fill lysosomal Ca^2+^ stores, the transient expression of TMEM165-mCherry in HEK293 human kidney epithelial cells increased the release of lysosomal Ca^2+^ upon ML-SA1-mediated activation of the lysosomal TRPML1 Ca^2+^ channel, as determined by the fluorescence intensity of the coexpressed TRPML1-GCaMP6s Ca^2+^ sensor (Extended Data Fig. 4a,b). Notably, the TMEM165-R126C mutant, which has been reported to increase the lysosomal localization of TMEM165 without affecting its activity (ref ^42^)^42^, increased the capacity of lysosomes to release Ca^2+^ in response to ML-SA1 even more than the wild-type TMEM165 (Extended Data Fig. 4a,b). Both the wild-type and the mutant TMEM165-mCherry colocalized with the lysosomal markers, LysoTracker and LAMP1, and their localization on the lysosomal limiting membrane was visualized in lysosomes enlarged by the PIKfyve inhibitor apilimod. (Extended Data Fig. 4c,d). Importantly, the transient expression of neither TMEM165 construct altered the expression levels of the TRPML1-GCaMP6s Ca^2+^ sensor (Extended Data Fig. 4e).

**Figure 4.**
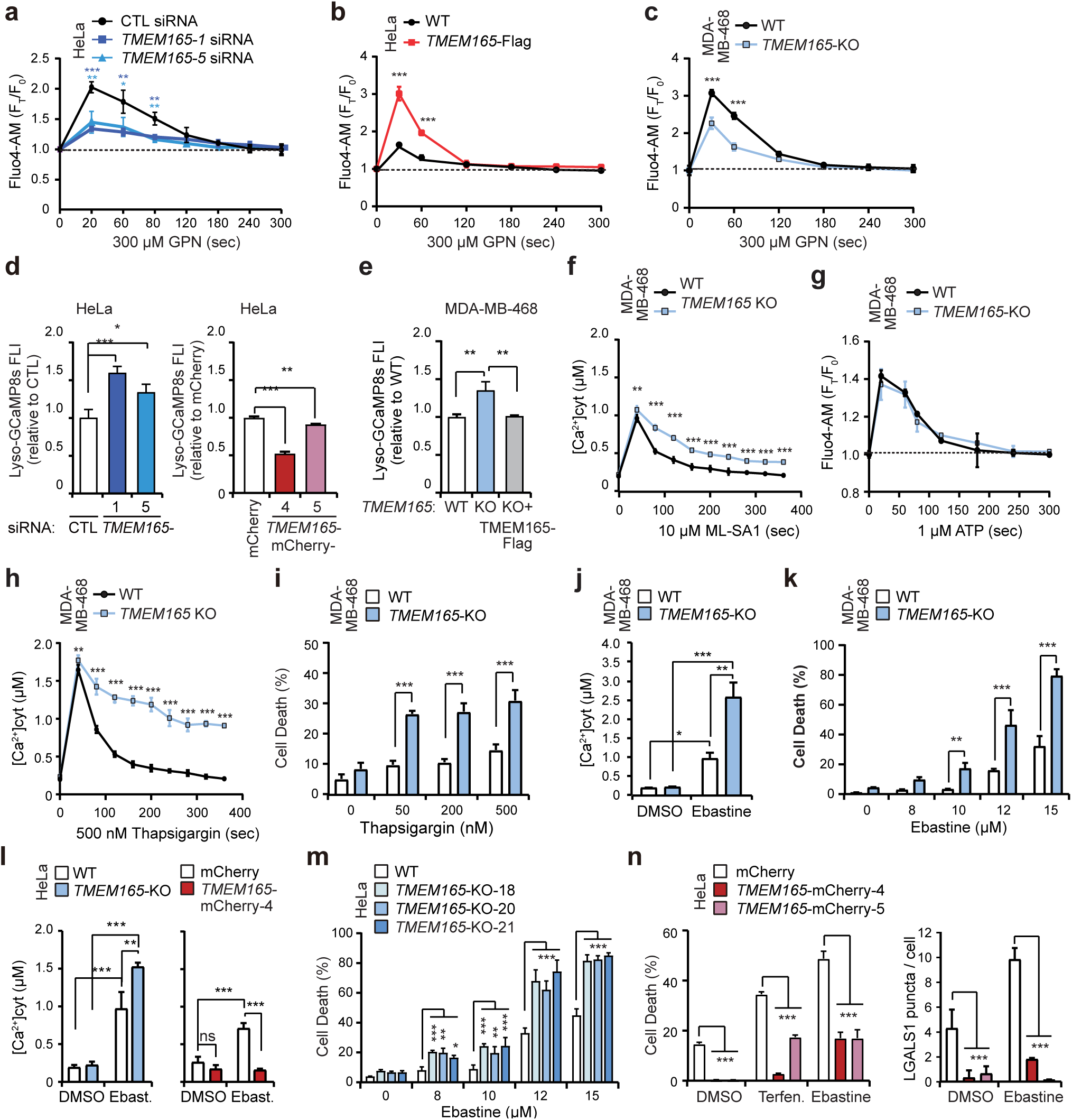
TMEM165 preserves lysosomal Ca2+stores and protects against cytosolic Ca2+ overload. a-c. Fluo-4-AM FLI in HeLa cells treated with indicated siRNAs for 72 h(a), HeLa TMEM165 overexpressed cells (b) and MDA-MB-468 TMEM165-KO cells (c) treated with 300 µM GPN for the last 0 – 300 sec. d. Lyso-GCaMP8s FLI in Hela cells treated for 72 h with CTL or TMEM165 siRNAs (left) or in HeLa TMEM165 overexpressed cells (right). e. Lyso-GCaMP8s FLI in indicated cell lines. f. Cytoslic calcium concentration analyzed by Fluo-4-AM FLI in WT and TMEM165-KO MDA-MB-468 clones treated with 10 µM MLSA1 for the last 0 – 300 sec. g. Fluo-4-AM FLI in WT and TMEM165-KO MDA-MB-468 clones treated with 1 µM ATP for the last 0 – 300 sec. h. Cytoslic calcium concentration analyzed by Fluo-4-AM FLI in WT and TMEM165-KO MDA-MB-468 clones treated with 500 nM thapsigargin for the last 0 – 300 sec. i. Death of indicated MDA-MB-468 cell clones treated with thapsigargin for 24 h. Cells were stained with propidium iodide (dead cells) and Hoechst-33342 (total cells) and cell death was analyzed by Celigo Imaging Cytometer. j. Cytoslic calcium concentration analyzed by Fluo-4-AM FLI in WT and TMEM165-KO MDA-MB-468 clones with 15 µM ebastine for 1 h. k. Death of indicated MDA-MB-468 cell clones treated with ebastine for 24 h. Cells were stained with propidium iodide (dead cells) and Hoechst-33342 (total cells) and cell death was analyzed by Celigo Imaging Cytometer. l. Cytosolic calcium concentration analyzed by Fluo-4-AM FLI in indicated WT and TMEM165-KO HeLa clones (left), mCherry-and TMEM165-mCherry-trasfected HeLa clones (right) with 15 µM ebastine for 1 h. m. Death of indicated HeLa cell clones treated with ebastine for 24 h. Cells were stained with propidium iodide (dead cells) and Hoechst-33342 (total cells) and cell death was analyzed by Celigo Imaging Cytometer. n. Death of indicated HeLa cell clones treated with terfenadine or ebastine for 24 h. Cells were stained with SYTOX Green (dead cells) and lysosomes) in mCherry- and TMEM165-mCherry-trasfected HeLa clones treated with 15 µM ebastine for the last 16 h (right). Error bars, SD of three independent experiments with ≥10 (f, h, j, l, n right), or ≥10000 (a, b, c, d, e, g, i, k, m, n left) randomly chosen cells analyzed in each sample. *, P < 0.05; **, P < 0.01; ***, P < 0.001 as analyzed by one-way Anova (d, e) or two-way Anova (a, b, c, f, g, h, i, j, k, l, m, n) with Tukey (d, e) or Dunnett (a, b, c, f, g, h, i, j, k, l, m, n) multiple comparison.

To study the perilysosomal free [Ca^2+^] without disturbing cellular Ca^2+^ homeostasis, we constructed a novel Ca^2+^ sensor, Lyso-GCaMP8s, by fusing the lysosomal membrane-targeting amino terminus of LAMTOR1 to the amino terminus of the GCaMP8s Ca^2+^ sensor. Supporting the role of TMEM165 in lysosomal Ca^2+^ influx, siRNA-mediated depletion of TMEM165 increased and TMEM165-mCherry expression reduced the perilysosomal free [Ca^2+^] in HeLa cells (Fig. 4d), and the reintroduction of TMEM165 into MDA-MB-468-TMEM165-KO cells completely reverted the high perilysosomal free [Ca^2+^] phenotype of the TMEM165-KO cells (Fig. 4e). Notably, TMEM165 depletion did not alter the free [Ca^2+^]_cyt_ in unstimulated MDA-MB-468 cells indicating that TMEM165 controls cellular Ca^2+^ homeostasis locally at the lysosomal membrane (Fig. 4f). Accordingly, TMEM165 deficiency delayed the clearance of cytosolic Ca^2+^ in MDA-MB-468 cells following ML-SA1-mediated activation of lysosomal TRPML1 Ca^2+^ channels but did not significantly affect the removal of cytosolic Ca^2+^ after ATP-induced activation of IP_3_ receptor-regulated Ca^2+^ channels in the ER (Fig. 4f,g). However, MDA-MB-468-TMEM165-KO cells failed to recover from the thapsigargin-induced cytosolic Ca^2+^ overload thereby revealing the potential of TMEM165 to clear cytosolic Ca^2+^ in the absence of the activity of ER Ca^2+^ pumps (Fig. 4h). Notably, the prolonged thapsigargin-induced cytosolic Ca^2+^ overload in TMEM165-deficient cells was associated with significantly increased cell death (Fig. 4i). Prompted by these results, we investigated the effect of TMEM165 on CAD-induced cancer cell death, which depends on lysosomal Ca^2+^ release and activation of Ca^2+^-dependent cyclic AMP synthesis ^27^. Compared to the parental MDA-MB-468 cells, TMEM165 deficiency significantly increased the ebastine-induced cytosolic Ca^2+^ overload, resulting in almost 3-fold higher free [Ca^2+^]_cyt_ 1 h after the treatment, and increased the subsequent cell death to a similar extent (Fig. 4j,k). TMEM165 deficient MDA-MB-468 cells displayed a similar sensitization to terfenadine (Extended Data Fig. 4f). The crucial role of TMEM165 in protecting cancer cells from CADs was confirmed in HeLa cells, where TMEM165 deficiency also increased the CAD-induced cytosolic Ca^2+^ overload and cell death while the expression of TMEM165-mCherry effectively inhibited both events (Fig. 4l-n). Notably, TMEM165-mCherry-mediated protection against CAD-induced HeLa cell death was associated with a nearly complete inhibition of CAD-induced formation of galectin 1 (LGALS1) puncta, indicative of the inhibition of lysosomal membrane permeabilization (Fig. 4n) ^45^.

Taken together, these data reveal taht TMEM165 is essential for maintaining lysosomal Ca^2+^ stores and protecting against cytosolic Ca^2+^ overload and subsequent lysosomal membrane permeabilization.

### A fraction of TMEM165 localizes to the lysosomal limiting membrane

TMEM165 was originally characterized as a Golgi resident protein, whose mutations cause congenital disorders of glycosylation ^39^. In line with the presence of the tyrosine-based, putative lysosomal targeting signal ^124^YNRL^127^ in the human TMEM165 protein, visualization of GFP- or RFP-tagged TMEM165 has revealed that TMEM165 traffics to lysosomes via the plasma membrane ^42^. To ensure that the TMEM165-mCherry used in this study also localized to lysosomal membranes, we analyzed its localization in HeLa and MDA-MBA-468 cells. In HeLa cells, the overexpressed TMEM165-mCherry formed strongly fluorescent punctuate structures colocalizing with the lysosomal markers Lyso-GFP (*r* = 0.93 ± 0.05) and Lysotracker^TM^ Green (*r* = 0.83 ± 0.05) and larger, perinuclear sheet-like structures with weaker fluorescence intensity and colocalization with the *trans-*Golgi marker EGFP-RAB6 (*r* = 0.42 ± 0.08) (Fig. 5a and Extended Data Fig. 5a). Indicating that the observed lysosomal localization was not caused by the overexpression, TMEM165-mCherry, which was expressed in MDA-MB-468-TMEM165-KO cells at a level similar to that of the endogenous TMEM165 protein, exhibited a similar distribution pattern with high intensity punctuate structures colocalizing with Lysotracker^TM^ Green (*r* = 0.85 ± 0.02) and sheet-like weaker intensity structures colocalizing with EGFP-RAB6 (*r* = 0.57 ± 0.12) (Extended Data Fig. 3e and 5b). Importantly, the formation of larger lysosomes upon inhibition of PIKfyve by vacuolin-1 revealed that also the stably expressed TMEM165-mCherry decorated lysosomal limiting membranes (Fig. 5a, *bottom*) ^46^, strongly suggesting that TMEM165-mCherry is a lysosomal membrane protein rather than a lysosomal cargo. This was further supported by the similar lysosomal localization of TMEM165-mCherry in cells in which autophagy was either pharmacologically (3-methyladenine) or genetically (ATG7-KO) inhibited (Fig. 5b).

**Figure 5.**
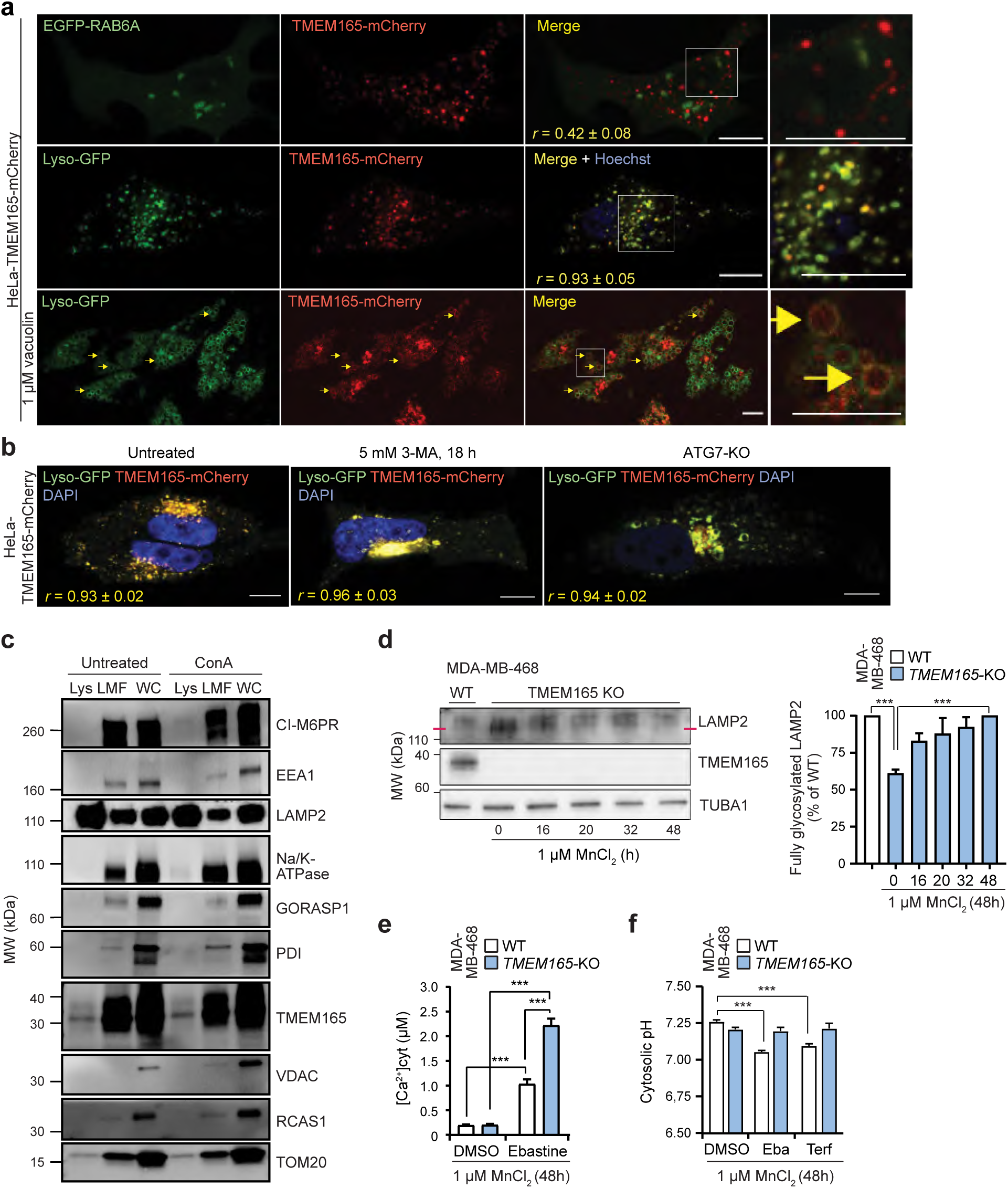
TMEM165 is present on the lysosomal limiting membrane. a. Representative (n = 3) confocal images of live HeLa-TMEM165-mCherry cells transfected with a Golgi marker EGFP-RAB6 (top), or stably expressing Lyso-GFP (middle and bottom). Yellow arrows point to vacuolin-enlarged lysosomes with both Lyso-GFP and TMEM165-mCherry on limiting membranes. DNA is visualized by Hoechst staining. White squares mark the area shown in enlarged images. r, Pearson’s co-localization coefficient (n=20). Scale bar, 10 µm. b. Representative (n=3) confocal images of live HeLa-TMEM165-mCherry-Lyso-EGFP cells with indicated treatments or genetic alteration. r, Pearson’s colocalization coefficient (n =20). Scale bars, 10 µm. c. Representative (n ≥ 3) immunoblots of indicated proteins from purified lysosomes (Lys), light membrane fractions (LMF) and whole cell lysates (WC) of HeLa cells. When indicated, 10 nM concanamycin A (ConA) was added to all buffers used for the extraction. d. Representative (n = 3) immunoblots of indicated proteins in WT and TMEM165-KO MDA-MB-468 clones treated with 1 µM MnCl2 for indicated times (left), and relative quantification of fully glycosylated LAMP2 (right). e. Cytoslic Ca2+ concentration analyzed by Fluo-4-AM FLI in WT and TMEM165-KO MDA-MB-468 clones treated with DMSO or 15 µM ebastine for the last 1 h of the 48 h treatment with 1 µM MnCl2. f. Cytosolic pH analyzed by pHrodo-AM FLI in WT and TMEM165-KO MDA-MB-468 clones treated with DMSO, 6 µM terfenadine, or 15 µM ebastine for the last 1 h of the 48 h treatment with 1 µM MnCl2. Error bars, SD of three independent experiments with ≥10 randomly chosen cells analyzed in each sample. *, P < 0.05; **, P < 0.01; ***, P < 0.001 as analyzed by by one-way Anova (d right) or two-way Anova (e, f) with Tukey (d right) or Dunnett (e, f) multiple comparison.

In line with previous reports ^39, 42^, immunostaining failed to reveal a significant colocalization of either endogenous or overexpressed TMEM165 with lysosomal markers (Extended Data Fig. 5c,d). To address this discrepancy, we realized that the localization of TMEM165-mCherry in HeLa cells dramatically altered from punctuate and mainly lysosomal structures in live or fixed cells to sheet-like and mainly Golgi-localized (colocalization with N-acetylgalactosaminyltransferase-EGFP (Golgi-EGFP, *trans*-Golgi)) structures following the permeabilization of fixed cells (Fig. 5a and Extended Data Fig. 5a,b,e). Due to this phenomenon, we were not able to demonstrate the lysosomal localization of the endogenous TMEM165 by immunostaining. Instead, we purified iron-dextran (FeDex)-loaded lysosomes from HeLa cells by magnetic capture and analyzed the proteins in the obtained eluates by immunoblotting. A small fraction of the endogenous TMEM165 was detected in the purified LAMP2-rich lysosomes, which did not contain detectable proteins from endosomes (early endosome antigen 1 (EEA1) and cation-independent mannose 6 phosphate receptor (CI-M6PR)), the Golgi (receptor-binding cancer antigen expressed on SiSo cells (RCAS1), Golgi reassembly stacking protein 1 (GORASP1), and CI-M6PR), the ER (protein disulfide isomerase (PDI)) or the plasma membrane (Na^+^/K^+^-ATPase) (Fig. 5c). However, the purified lysosomal fraction contained a small amount of mitochondrial import receptor subunit TOM20 homolog (TOM20) possibly due to copurified mitochondria attached to lysosomes or mitophagy (Fig. 6c). Notably, the inhibition of the V-ATPase by concanamycin A during the lysosome purification procedure, which inhibits the degradation of cargo proteins in the lysosomal lumen, did not significantly alter the level of TMEM165 in purified lysosomes, while the putative cargo proteins Na^+^/K^+^-ATPase and CI-M6PR became detectable after concanamycin A treatment (Fig. 5c).

**Figure 6.**
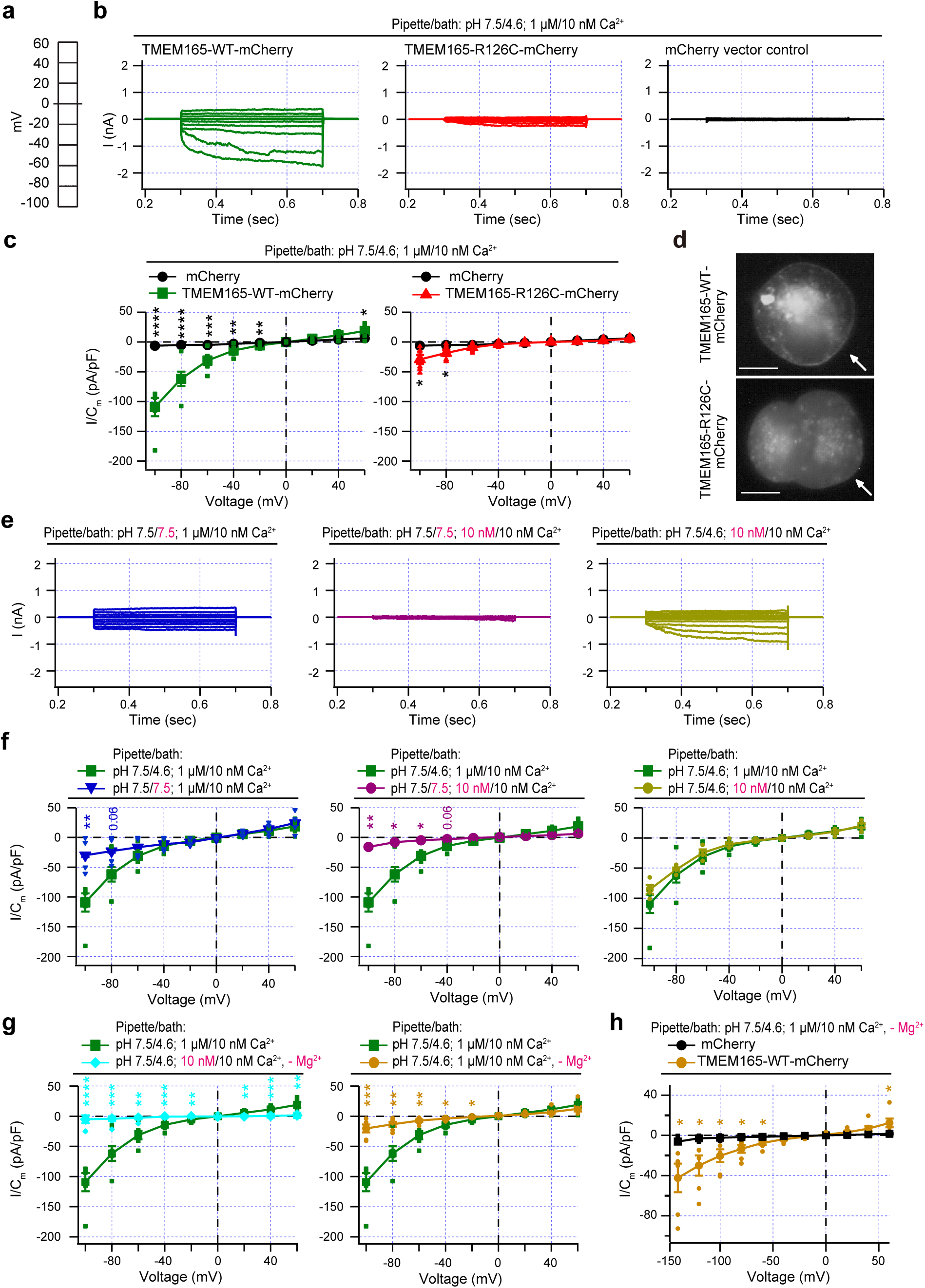
TMEM165 mediates H+ inward and Ca2+ outward currents. **a**. Voltage step protocol employed for current recordings. **b**. Examples of macroscopic current recordings conducted for TMEM165-WT-mCherry (green), TMEM165-R126C-mCherry mutant (R126C, red), and mCherry control (black) transiently expressed in HEK293 cells. Measurements were performed using indicated pH and Ca2+ concentrations in the pipette/bath. **c**. Current densities plotted against respective voltages for TMEM165-WT-mCherry (left, green squares, n=6) and TMEM165-R126C-mCherry (right, red triangles, n=6) and the control (black circles, n=12). **d**. Representative live images of HEK293 cells transiently transfected with TMEM165-WT-mCherry or TMEM165-R126C-mCherry. Arrows, plasma membrane; scale bars, 10 µm. **e**. Examples of macroscopic current recordings for TMEM165-WT-mCherry using indicated pH and Ca2+ concentrations in the pipette/ bath. **f**. Current densities plotted against respective voltages for TMEM165-WT-mCherry at indicated pH and Ca2+ concentrations in the pipette/bath solution with asymmetrical Ca2+ and pH (green squares, n=6) or asymmetrical Ca2+ and symmetrical pH (blue reverted triangles, n=4), symmetrical Ca2+ and symmetrical pH (magenta circles, n=3), and symmetrical Ca2+ and asymmetrical pH (mustard circles, n=4). **g**. Current densities plotted against respective voltages for TMEM165-WT-mCherry at indicated pH and Ca2+ concentrations in the pipette/bath solution with asymmetrical Ca2+ and pH with Mg2+ (green squares, n=6) and without Mg2+ (mustard circles, n=6, right) or symmetrical Ca2+ and asymmetrical pH without Mg2+ (cyan circles n=7, left). **h**. Current densities plotted against respective voltages for TMEM165-WT-mCherry (orange circles, n=6) and mCherry control (black squares n=6) at indicated pH and Ca2+ concentrations in the pipette/bath solution with asymmetrical Ca2+ and pH, and without Mg2+. Error bars, ± SEM; *, P < 0.05; **, P < 0.01; ***, P < 0.001, ****, P < 0.0001, or as analyzed by unpaired t-test using GraphPad Prism 10.0.2.

Because the TMEM165-mediated maintenance of the Golgi Mn^2+^ homeostasis is essential for the proper N-glycosylation of several membrane proteins ^40^, we next asked whether the lysosomal phenotype observed in TMEM165 depleted cells could be a result of the defective glycosylation of the normally highly lysosomal membrane proteins. Treatment with 1 µM MnCl_2_ for 48 h completely reversed the hypoglycosylation phenotype in MDA-MB-468-TMEM165-KO cells as demonstrated by the MnCl_2_-induced reversal of the increased mobility of LAMP2 in sodium dodecyl sulfate–polyacrylamide gel electrophoresis (Fig. 5d). However, the MnCl_2_ supplementation rescued neither the prolonged cytosolic Ca^2+^ peak nor the defective cytosolic acidification in CAD treated MDA-MB-468-TMEM165-KO cells (Fig. 5e,f).

These data support the idea that a small fraction of TMEM165 localizes to the lysosomal limiting membrane where it regulates lysosomal ion homeostasis.

### TMEM165 mediates H^+^ and Ca^2+^ dependent currents

The data presented above strongly suggest that TMEM165 plays a role in Ca^2+^ influx to the lysosome, and H^+^ efflux from the lysosome to the cytosol. To test this hypothesis, we performed patch-clamp experiments using HEK293 cells transfected with wild-type TMEM165-mCherry, the R126C mutant, or the mCherry vector as a control. We applied a step protocol from −100 mV to +60 mV with 20 mV increments in the whole-cell mode with an asymmetrical pH (pipette 7.5/bath 4.6) and [Ca^2+^] (pipette 1 µM/bath 10 nM) across the membrane mimicking the conditions in the cytosol (pipette) and the lysosomal lumen (bath) (Fig. 6a). In cells expressing wild-type TMEM165-mCherry, we detected significant inward current activity (most likely the net current of H^+^ inward and Ca^2+^ outward) in a physiologically relevant voltage range that was absent in control cells (Fig. 6b,c). TMEM165-R126C mutant currents were considerably weaker than wild-type currents (Fig. 6b,c), while in the lysosomal Ca^2+^ imaging experiments, the mutant protein had stronger effect on Ca^2+^ release than the wild-type protein (Extended Data Fig. 4a,b). This difference is presumably due to the increased lysosomal and decreased plasma membrane localization of the mutant protein (Fig. 6d; Extended Data Fig. 4c,d) (ref ^42^). Thus, we focused on wild-type TMEM165-mCherry and increased the bath pH from 4.6 to 7.5 with the result that the measured inward current was almost completely reduced to zero (Fig. 6e,f, left). Additionally, we reduced the Ca^2+^ concentration in the pipette from 1 µM to 10 nM, resulting in a similar, almost complete loss of the current (Fig. 6e,f, middle). However, when omitting only Ca^2+^ we noticed a comparably small reduction in the current (Fig. 6e,f, right), letting us to speculate that this may be the result of the presence of 1 mM Mg^2+^. We therefore next omitted both Ca^2+^ and Mg^2+^ from the bath, now resulting in a complete loss of the current (Fig. 6g, left). When only Mg^2+^ was omitted, the current decreased less (Fig. 6g, right) and remained significant in the TMEM165-expressing cells when compared to control cells (Fig. 6h). These data suggest that the omission of either Ca^2+^ (in the absence of Mg^2+^) or H^+^ significantly affects the observed current.

Although PIKfyve inhibitors enlarged TMEM-165-mCherry-positive lysosomes to some degree (Fig. 5g; Extended Data Fig. 4d), our efforts to detect TMEM165 currents across the lysosomal limiting membrane were unfortunately hindered by the failure of PIKfyve inhibitors to trigger the formation of large enough TMEM165-positive giant lysosomes for patch clamp, possibly due to TMEM165-induced disturbances in the lysosomal Ca^2+^ homeostasis.

These data indicate that TMEM165 expression results in the voltage-dependent development of H^+^ and Ca^2+^ dependent currents, which are not present in controls. Thus, TMEM165 is indeed capable of transporting Ca^2+^ and Mg^2+^ ions into lysosomes and H^+^ ions out of lysosomes in an antiporter like manner.

## Discussion

Lysosomal fitness depends on tightly regulated pH and Ca^2+^ concentration gradients across the limiting membrane ^2, 14^. While a low pH is essential for the activity of most lysosomal hydrolases, the ligand-activated Ca^2+^ release through TRPML1, two pore channels, and P2RX4 governs endolysosomal membrane trafficking and initiates numerous signaling cascades controlling for example lysosomal biogenesis, cellular metabolism and lysosomal membrane stability ^27, 47–49^.

Despite the crucial role of lysosomal Ca^2+^ stores in the maintenance of lysosomal and cellular health, the identity of the protein(s) responsible for filling mammalian lysosomes with Ca^2+^ remains unknown. The data presented above identifies TMEM165 not only as a protein responsible for maintaining lysosomal Ca^2+^ stores but also as a mediator of Ca^2+^-induced lysosomal H^+^ leakage in mammalian cells. The role of TMEM165 in maintaining lysosomal Ca^2+^ stores is supported by ample data showing i) a significant reduction in the amount of Ca^2+^ released to the cytosol upon GPN-mediated lysosomal membrane permeabilization in TMEM165-depleted HeLa and MDA-MB-468 cells; ii) increased capacity of lysosomes to release Ca^2+^ in TMEM165 overexpressing HeLa and HEK293-TRPLM1 cells upon treatment with GPN and ML-SA1, respectively; iii) TMEM165-dependent recovery from the cytosolic Ca^2+^ overload in thapsigargin-, ML-SA1- and ebastine-treated MDA-MB-468 cells and ebastine-treated HeLa cells; and finally, iv) the development of voltage-dependent Ca^2+^ currents observed when patch-clamping TMEM165 overexpressing HEK293 cells. Previous studies have demonstrated the dependence of lysosomal Ca^2+^ influx on ER Ca^2+^ stores, ER-lysosome contact sites and acidic lysosomal pH ^9, 50–52^. The reported delay in the inhibition of lysosomal Ca^2+^ influx following the disruption of the lysosomal pH gradient has, however, suggested that the pH dependence of lysosomal Ca^2+^ influx observed in living cells is caused by the disruption of ER-lysosome contact sites rather than the loss of the lysosomal pH gradient ^53^. Based on our data, TMEM165-mediated Ca^2+^ influx is however highly dependent on the pH gradient, whose removal in our patch-clamp experiments abolished the TMEM165-mediated currents almost completely. This is further supported by a recent manuscript by Dr. Krishnan and coworkers, which introduced TMEM165 as a proton-activated lysosomal Ca^2+^ importer ^54^.

In addition to demonstrating the ability of TMEM165 to import Ca^2+^ from the cytosol to the lysosome, our data also highlight the ability of TMEM165 to protect mammalian cells against Ca^2+^ overload. This finding is in line with the reported sensitivity of the yeast strain depleted of the TMEM165 ortholog Gdt1p (*gdt1*Δ) to high external Ca^2+^ concentrations, and the ability of the amino-terminally truncated human TMEM165 to partially restore the growth of the *gdt1*Δ strain under Ca^2+^ stress ^28^. While the mammalian TMEM165 may play only a moderate role in the clearance of cytosolic Ca^2+^ in general, its role in removing Ca^2+^ locally from the perilysosomal area is likely to be of greater importance. This is clearly demonstrated by the substantial sensitization to and inhibition of CAD-induced lysosome-dependent cancer cell death upon TMEM165 depletion and overexpression, respectively. These data emphasize the role of P2RX4-mediated lysosomal Ca^2+^ release in CAD-induced cancer cell death and are in line with our previous identification of Ca^2+^-dependent adenylate cyclase 1 as a mediator of CAD-induced lysosomal membrane permeabilization and cell death ^27, 34^. Providing further support for a link between the lysosomal P2RX4 Ca^2+^ channel and TMEM165-mediated lysosomal H^+^ leakage, a recent report describes a significant, P2RX4-dependent increase in the lysosomal pH in metastatic breast cancer cells ^55^, and the expression of both P2RX4 and TMEM165 has been linked to the metastatic potential of breast cancer cells ^55, 56^.

To study the molecular basis of the lysosomal H^+^ leakage conductance, we created a fluorescent pH sensor, Lyso-mKeima, that detects the local pH on the surface of lysosomes. In the meanwhile, Haoxing Xu and colleagues identified TMEM175 as the long-sought channel responsible for the H^+^-activated lysosomal H^+^ leakage, which maintains a lysosomal pH of approximately 4.5 ^17^. Using cytosolic pH indicators and Lyso-mKeima to measure cytosolic and perilysosomal pH, respectively, we demonstrated a substantial TMEM175-dependent acidification of the perilysosomal area in unstimulated HeLa cells and a further, TMEM175-independent acidification in response to Ca^2+^-mobilizing agents. The latter was effectively inhibited by the chelation of cytosolic free Ca^2+^ by BAPTA-AM and by depletion of TMEM165, strongly suggesting that the TMEM165-mediated lysosomal Ca^2+^ uptake triggers a TMEM165-dependent lysosomal H^+^ leakage. This hypothesis is strongly supported by our electrophysiological experiments demonstrating the ability of TMEM165 to mediate voltage-dependent H^+^ and Ca^2+^ dependent currents. The observed low pH on the lysosomal surface in unstimulated cells, and especially the further, TMEM165-dependent long-lasting acidification induced by Ca^2+^ mobilizing agents is highly intriguing and argues against the idea that variations in the cytosolic pH are insignificant, because cellular buffers quickly stabilize them. Such drops in pH are known to alter the protonation status and function of several proteins, for example phosphofructokinase, whose glycolytic activity regulates AMP-dependent protein kinase (AMPK) on the lysosomal membrane ^57, 58^, and the GTP-binding protein RHEB, which controls the activity of mammalian target of rapamycin complex 1 (mTORC1) by recruiting it to the lysosomal membrane ^59^. Thus, it is tempting to speculate that Ca^2+^-induced acidification of the perilysosomal area could contribute to the Ca^2+^ overload-induced shift from anabolic to catabolic metabolism by altering the activity of AMPK, mTORC1 or other metabolic regulators residing on the lysosomal membrane ^12, 58, 60^.

As discussed above, the maintenance of lysosomal H^+^ and Ca^2+^ homeostasis is essential for cellular fitness. Accordingly, their disturbances are associated with a wide range of pathologies ranging from degenerative disorders to cancer ^3, 4, 12, 16, 20, 61^. Thus, the identification of TMEM165 as a regulator of lysosomal H^+^ efflux and Ca^2+^ influx opens new avenues for the development of therapeutics for the treatment of these diseases. In particular, TMEM165 antagonists and their combination with CADs or other Ca^2+^ mobilizing cancer drugs could prove highly effective in the future cancer treatment by increasing and prolonging the cytosolic Ca^2+^ overload and promoting the subsequent lysosomal membrane permeabilization, while TMEM165 agonists could reverse defective lysosomal Ca^2+^ stores in lysosomal storage disorders and assist the recovery of neurons from the cytosolic Ca^2+^ overload in Parkinson’s disease.

## Materials and Methods

### Cell lines

HeLa cervical, human, female carcinoma cells were obtained from the European Collection of Authenticated Cell Cultures (ECACC, 93021013). The TNF-sensitive S1 subclone of human, MCF7 female breast cancer cells has been described previously ^62^. MDA-MB-231 (HTB-26) and MDA-MB-468 (HTB-132) human, female breast cancer cell lines, MCF10A (CRL-10317) human female breast epithelial cells, and human kidney HEK-293 (CRL-1573) cells originating from a female fetus were obtained from the American Type Culture Collection (ATCC). TMEM165-KO cell lines were generated using CRISPR/Cas9-mediated gene editing by transfecting single-guide RNA targeting TMEM165 (PXPR001TMEM165) into HeLa and MDA-MB-468 cells, and after 2 weeks clones were picked up and validated by immunoblotting and sequencing. HeLa cells were cultured in Dulbecco’s modified Eagle medium (DMEM; Thermo Fisher Scientific, 31966021) supplemented with 10% heat-inactivated fetal calf serum (Thermo Fisher Scientific, 10270-106) and penicillin/ streptomycin (Gibco, 15140122). MDA-MB-231 and MDA-MB-468 cells were cultured in DMEM supplemented with 2mM Glutamine, 1x MEM nonessential amino acids solution (Thermo Fisher Scientific, 11140035), 10% heat-inactivated fetal calf serum and penicillin/ streptomycin. MCF-7 cells were cultured in RPMI 1640 (Thermo Fisher Scientific, 61870010) supplemented with 10% heat-inactivated fetal calf serum and penicillin/ streptomycin. MCF-10A cells were cultured in DMEM/ F12 (Thermo Fisher Scientific, 31330038) supplemented with 5% horse serum (Thermo Fisher Scientific, 16050-122), 20 ng/ ml epidermal growth factor (Sigma-Aldrich, e-4127), 0.5 µg/ ml hydrocortisone (Sigma-Aldrich, 3867), 100 ng/ ml choleratoxin (Sigma-Aldrich, c-8052), 10 µg/ ml insulin (Sigma-Aldrich, I9278) and penicillin/ streptomycin. HEK293 cells were cultured in DMEM supplemented with penicillin/ streptomycin and 10% fetal bovine serum. All cells were cultured at 37°C in a humidified chamber with 95% air and 5% CO_2_ and tested regularly for mycoplasma.

### Plasmids

The Lyso-mKeima expression construct was generated by replacing Lyso-GFP in pLV-Lyso-GFP-puro (kindly provided by Dr. Shawn M. Ferguson, Yale University, New Haven, CT) with the first 39 amino acids of LAMTOR1 and the GGSGGS linker (Lyso) using restriction sites EcoRI and BamHI. Then the mKeima was inserted into pLV-Lyso-puro using the restriction sites BamHI and XbaI. The TMEM165-mCherry expression construct was generated by inserting mCherry into pLentiCMVie-IRES-BlastR (Addgene #119863) using the restriction sites XhoI and BamHI. Then, the TMEM165 cDNA was amplified from pCDNA3.1NEGFP-TMEM165 (Genscript) and inserted into pLentiCMVie-mCherry-BlastR via NheI and XhoI sites. The Lyso-GCaMP8s expression construct was generated by inserting Lyso into pLentiCMVie-IRES-BlastR (Addgene #119863) via the EcoRI and EcoRV sites. Then, the GCaMP8s was amplified from pGP-CMV-jGCaMP8s (Addgene #162371) and inserted into pLentiCMVie-lyso-BlastR using the restriction sites EcoRV and BamHI. The TMEM165-flag expression construct was generated by inserting TMEM165-flag into pLentiCMVie-IRES-BlastR via the NheI and XhoI sites. The TRPML1-GCamP6s expression construct was created by subcloning the human TRPML1 insert from the human TRPML1-EYFP plasmid ^63^ into the pGP backbone harboring GCaMP6s (Addgene #40753) using restriction sites BglII (N-terminal) and SalI (C-terminal).

PXPR001TMEM165 was generated by using the gRNA: GCAGC CGGGG CGCCG AUGCG. Golgi-EGFP (#79809), EGFP-RAB6A (#49469) and LAMP1-mGFP (#34831) were purchased from Addgene. pcDNA3.1+-mCherry was kindly provided by Dr. Christian Wahl-Schott (Biomedical Center, LMU, Munich, Germany).

**Table 1.**
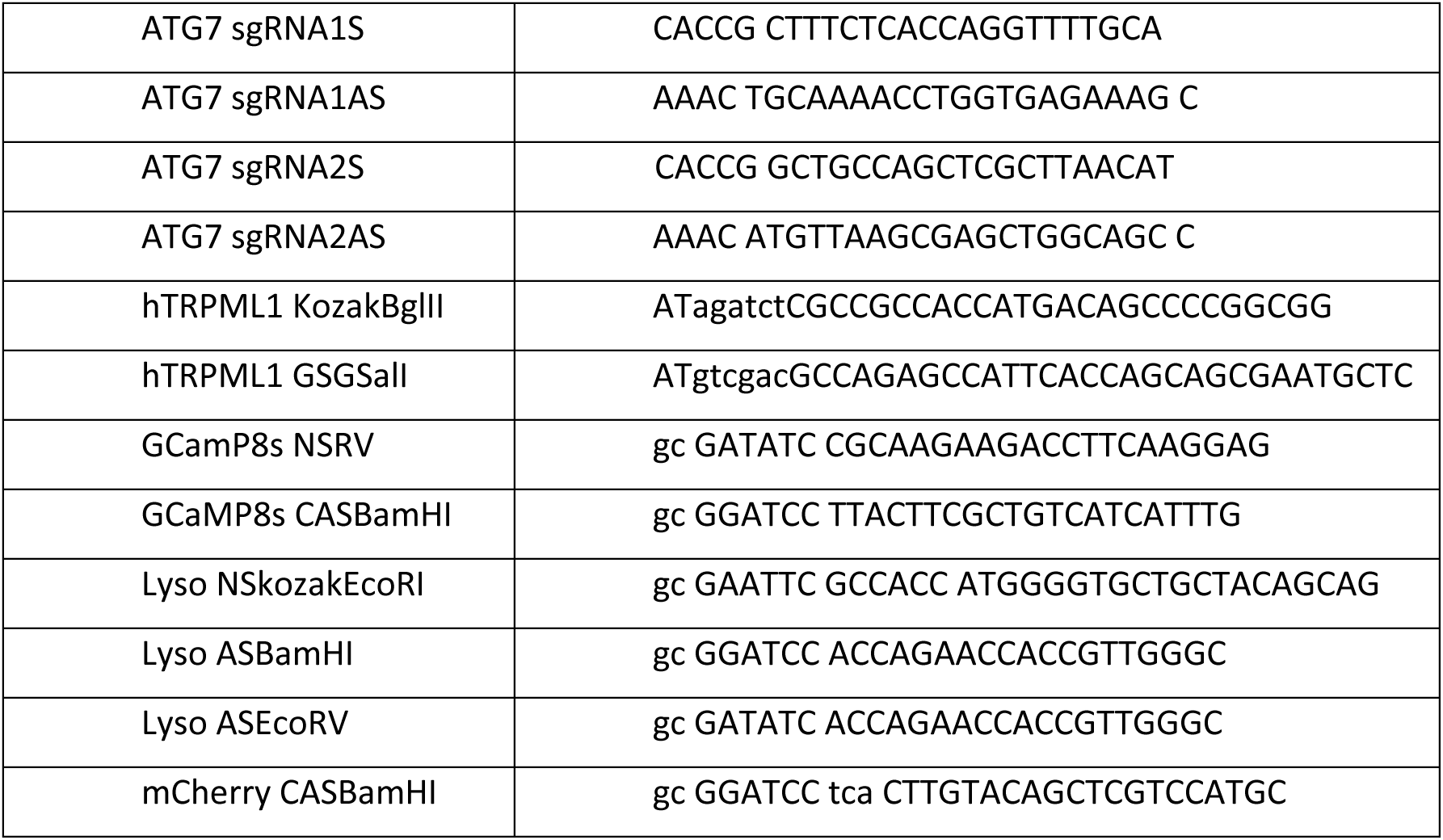
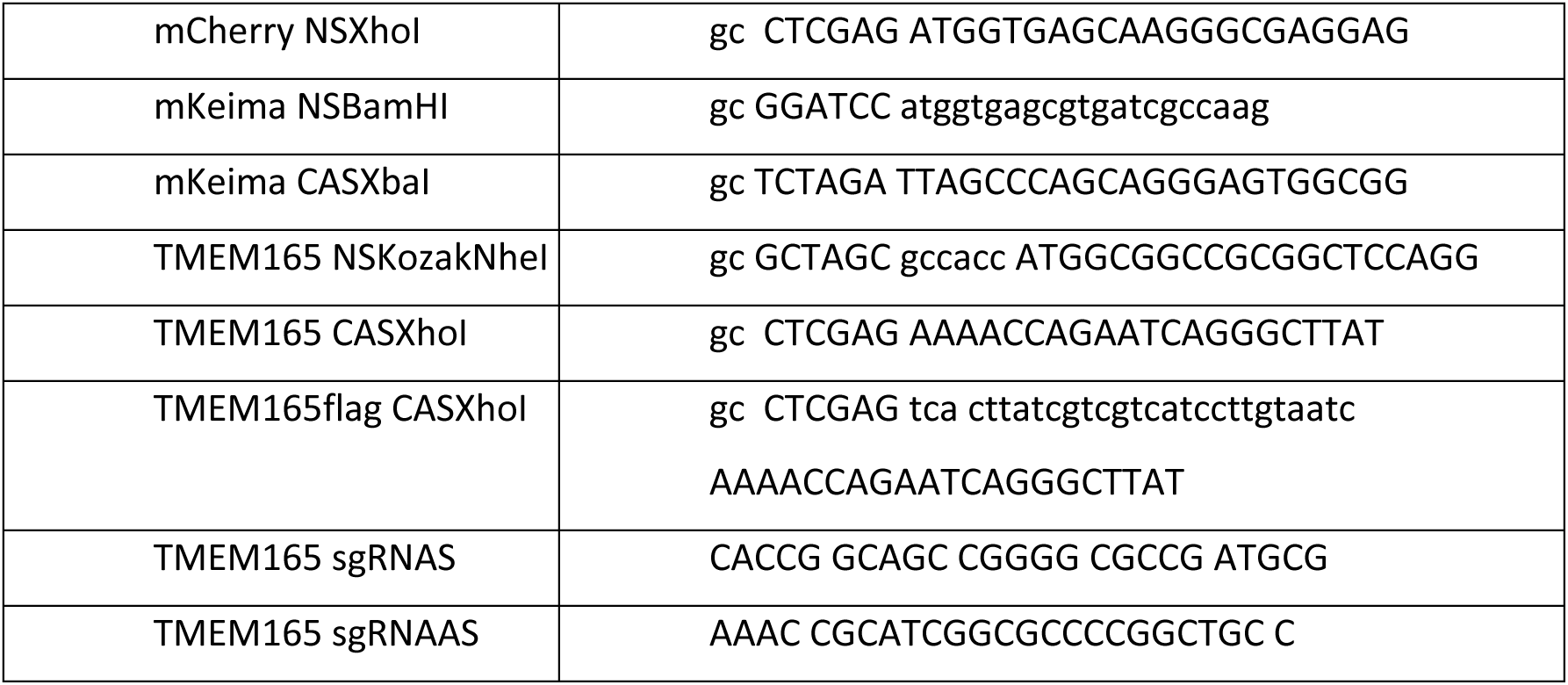
Cloning primers.

### siRNAs

P2X4: CAAGUCGUGCAUUUAUGAUtt/AUCAUAAAUGCACGACUUGtt, GUCCUCUACUGCAUGAAGAtt/UCUUCAUGCAGUAGAGGACtt. TMEM165: GUAUCUGAAUUG GGUGAUAtt/UAUCACCCAAUUCAGAUACaa,CAGGGUCUAUACAUACUAUtt/AUAGUAUGUAUAGACCC UGaa. TMEM175: GCCUUACACGUUUUCGUUAtt/ UAACGAAAACGUGUAAGGCag, CUGCA UGAUG ACCAU CACC aa/GGTG AUGGT CAUCA UGCAGag.

### Antibodies

Anti-α-Tubulin (ab40742), anti-β-Actin (ab20272), anti-GFP (Ab290), anti-Lamin B1 (ab194109), anti-Galectin 1 (ab25138), anti-P4HB (ab137110), anti-EEA1 (ab2900), anti-sodium potassium ATPase (ab76020), and anti-M6PR (ab124767) were purchased from Abcam; anti-GAPDH (5174S), anti-RCAS1 (12290), and anti-VDAC (12454S) from Cell Signaling Technology; anti-LAMP2 (L0668-200UL) from Sigma Aldrich; anti-LAMP1 (sc20011) and anti-Tom20 (sc-11415) from Santa Cruz; anti-TMEM175 (PA5-100355), anti-mouse IgG Alexa Fluor 488 (A-11001), anti-rabbit IgG Alexa Fluor 594 (A-21207), and anti-rabbit IgG Alexa Fluor 488 (A-21206) from Thermo Fisher Scientific; anti-TMEM165 (20485-1-AP) from Proteintech Europe; anti-P2RX4 (APR-002) from Labome; and goat anti-rabbit IgG (PI-1000) from Vector Laboratories.

### Chemicals and reagents

Ammonium hydroxide solution (28.0-30.0%) (221228), D-(+)-glucose (G7021), Hoechst-33342 (B2261), iron (II) chloride (372870), iron (III) chloride (157740), NMDG (66930), propidium iodide (P4864), terfenadine (T9652), and valinomycin (V0627) were purchased from Sigma Aldrich. BAPTA-AM (B1205), CellMask™ Green (C37608), ER-Tracker™ Green (E34251), Fluo-4-AM (F14201), HEPES (BP310), LysoTracker™ Deep Red (L12492), LysoTracker™ Green DND-26 (L7526), MitoTracker™ Green (M7514), Nigericin (N1495), and SYTOX™ Green, Nucleic Acid Stain (S7020) were obtained from Thermo Fisher Scientific. ML-SA1 (4746) and thapsigarin (1138) were from Tocris, ebastine (15372) from Caymann, arachidonic acid (HY-109590) from MedchemExpress, concanamycin A (sc-202111) from Santa Cruz, dextran 40 BioChemica (A2249) from PanReac AppliChem, hydrochloric acid (VWRC30024.290) from VWR, MES (02195309-CF) from MP Biomedicals, and EGTA (324626) from Merck.

### Transfections

Plasmid transfections were performed by using TurboFectin 8.0 (Origene, TF81001), Lipofectamine 3000 transfection agent (Thermo Fisher Scientific, L3000008) or Turbofect^TM^ (Thermo Fisher, R0531) according to the manufacturer’s instructions. siRNA transfections were performed by using Lipofectamine RNAiMax (Thermo Fisher Scientific, 13778075) according to the manufacturer’s instructions.

### Western blot

Cell lysates were prepared in Laemmli sample buffer (125 mM Tris, pH 6.7; 20% glycerol; 140mM SDS) supplemented with complete protease inhibitor cocktail (Roche, 04693159001) and PhosSTOP™ (Roche, 04906837001), and 0.05 M dithiothreitol and then loaded onto a 4%–20% gradient SDS-polyacrylamide gel for electrophoresis. Proteins were transferred onto polyvinylidene difluoride membranes by using a Bio-Rad Trans-Blot Turbo system. The membranes were blocked with PBS containing 5% milk and 0.1% Tween-20 and incubated with the indicated primary antibodies and corresponding peroxidase-conjugated secondary antibodies. The signal was detected with Clarity Western ECL Substrate and Luminescent Image Reader, and quantified by densitometry with Image Studio Lite software.

### Immunostaining

Cells were grown on coverslips and fixed in 4% paraformaldehyde in Dulbecco phosphate-buffered saline (DPBS; Thermo Fisher Scientific, 14190094) for 20 min were permeabilized with 0.1% saponin in DPBS for 10 min and blocked in 5% goat serum in DPBS for 10 min. Sequential incubation of the cells with the indicated primary antibodies and corresponding Alexa Fluor 488- or Alexa Fluor 594-coupled secondary antibodies was performed to label the protein of interest. Nuclei were labeled either with 5 mg/ml Hoechst-33342 or DAPI in the Prolong Gold Antifade Mounting medium. Unless otherwise indicated, images were acquired with an LSM700 microscope with a Plan-Apochromat 63×/1.40 Oil DIC M27 objective and Zen 2010 software (all equipment and software from Carl Zeiss, Jena, Germany) and analyzed with Image J (Fiji) software.

For live imaging, cells were treated for 10 min with CellMask^TM^ Green dye (1:1000) at 37°C, then treated with LysoTracker^TM^ DeepRed for 30 min at 37°C, washed three times in PBS, treated with a Hoechst-33342 and then processed for live imaging with an Opera Phenix High Content Confocal Imaging System (Revvity).

### Ca^2+^ imaging

For imaging with Fluo-4-AM, ells were stained with 3 mM Fluo-4-AM for 25 minutes in DMEM (Thermo Fisher, 10567014). After two washes with DPBS, the cells were resuspended in Dulbecco’s PBS (Gibco, 14190-094) plus 20 mM HEPES. Further incubation with the indicated treatments was performed at 37°C. Relative quantification and comparison of Fluo-4-AM fluorescence were performed using a BD FACSVerse flow cytometer.

To measure the absolute [Ca^2+^]_cyt_, cells were grown in 96-well plates and stained with 3 mM Fluo-4-AM in medium for 25 min. After the indicated treatments, we changed the medium to HBSS (Thermo Fisher Scientific, 14025050) and imaged the cells on the ImageXpress Pico platform. The absolute [Ca^2+^]_cyt_ was calculated according to

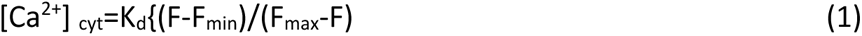

F is the desired fluorescence intensity measured by the ImageXpress Pico platform; F_max_ is the fluorescence intensity at saturating free Ca^2+^ concentration ([Ca^2+^] _free_); F_min_ is the fluorescence intensity at zero [Ca^2+^] _free_; To determine K_d,_ we stained the cells with 3 mM Fluo-4-AM and then incubated the cells with a series of Ca^2+^ calibration buffers ([Ca^2+^] _free_ =0, 0.0717, 0.1613, and 39.8 μM) supplemented with 10 µM valinomycin, 10 µM nigericin and 3 µM ionomycin for 5 min. Thereafter, the images of the cells were taken with the ImageXpress Pico platform. The obtained fluorescence intensities of Fluo4-AM were used to plot log {(F-F_min_)/(F_max_-F)} vs. Log [Ca^2+^]. The X-intercept from the linear plot is the Log K_d_. For the preparation of calcium calibration buffer, the [Ca^2+^] _free_ was estimated using the equation:

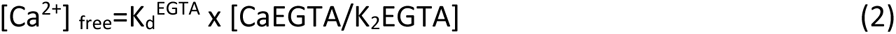

where K_d_^EGTA^ is the dissociation constant of CaEGTA at a given temperature, ionic strength and pH and [CaEGTA/K_2_EGTA] is the molar ratio of CaEGTA to K_2_EGTA in the particular solution. The table listing K_d_^EGTA^ values for CaEGTA in 0.1 M KCl at 37°C can be found through the link (https://biotium.com/wp-content/uploads/2013/07/PI-59100.pdf).

HEK293 cells were seeded on 25 mm^2^ glass coverslips in standard tissue culture plates (6-well plate, 2 mL of total volume per well) at a density of 2×10^5^ cells per well, i.e., 100,000 cells per mL. After 48 h in culture, the cells were transfected. HEK293 cells were transfected with 1.5 μg of TRPML1-GCaMP6s- or 1 μg of the respective control plasmid or TMEM165-mCherry-carrying plasmid, 4 μL of Turbofect^TM^, and 200 μl of serum-free DMEM per well in a 6-well plate. The cells were then incubated at 37°C for 24 h. Ca^2+^ imaging was performed using an inverted Leica DMi8 live cell microscope. First, transfected cells were washed with DMEM and Ca^2+^-free buffer was used to carefully rinse the wells before placing the glass coverslips into the imaging chamber. All GCaMP6s experiments were conducted in Ca^2+^-free buffer comprising 138 mM NaCl, 6 mM KCl, 1 mM MgCl_2_, 10 mM HEPES, 5.5 mM D-glucose monohydrate and 2 mM EGTA (adjusted to pH = 7.4 with NaOH). Then, 450 μl Ca^2+^-free buffer was slowly added to the chamber to prevent the cells being washed away. The osmolarity of the Ca^2+^-free buffer was adjusted to 300 mOsmol/L. GCaMP6s was excited at 470 nm (GFP excitation wavelength) and the emitted fluorescence was captured with a 515 nm longpass filter. mCherry was excited at 568 nm and the emitted fluorescence was captured with a 590 nm filter. Images were obtained every 2.671 sec with a 63x objective. Regions of interest were drawn around each cell coexpressing the GCaMP6s and mCherry constructs. Background areas without cells were selected for manual background subtraction. The fluorescence intensity was calculated using LAS X 5.1.0 software. Baseline values were acquired by averaging fluorescence intensity values from a 30 s recording before the addition of 10 μM ML-SA1. The area under the curve was calculated using GraphPad Prism 9.0.1 software. The mean values per experiment containing 4-12 cells were calculated and used for plotting.

### Perilysosomal pH measurement

Lyso-mKeima expressing cells were grown in 96-well plate and then treated as indicated. Finally, we changed the medium to Live Cell Imaging Solution (Thermo Fisher Scientific, A14291DJ) and obtained images of the cells with the ImageXpress high-content platform. Standard curves used to estimate perilysosomal pH were created by a similar analysis of cells incubated with a series of pH calibration buffers (pH 6.0, 6.5, 7.0, and 7.5; 140 mM NaCl; 2.5 mM KCl; 1.8 mM CaCl_2_; 1mM MgCl_2_; and 20 mM HEPES) supplemented with 10 µM valinomycin and 10 µM nigericin for 5 min.

### Lysosomal pH measurement

To calculate the absolute lysosomal pH, we plated cells in 96-well plates. The next day, the cells were stained with 1 μM LysoSensor Yellow /Blue DND-160 (Thermo Fisher Scientific, L-7545) for 5 minutes, washed twice with DPBS. Then we changed the medium to phenol-red-free DMEM supplemented with high glucose and HEPES (Thermo Fisher Scientific, 21063029) and measured the fluorescence of the cells in the SpectraMax iD3 at Ex 329 nm/ Em 440 nm and Ex 380 nm/ Em 540 nm. Standard curves used to estimate lysosomal pH were created by a similar analysis of cells incubated with a series of pH calibration buffers (pH 4, 4.5, 5, and 5.5, 15 mM HEPES, 130 mM KCl, 1 mM MgCl_2_) supplemented with 10 µM valinomycin and 10 µM nigericin for 5 min.

### Cytosolic pH measurement

Stably transfected HeLa SypHer3S cells were grown on 96-well plates. After the indicated treatments, the medium was changes to Live Cell Imaging Solution and images were taken with the ImageXpress high-content platform. Standard curves used to estimate cytosolic pH were created by a similar analysis of cells incubated with a series of pH calibration buffers (pH 6, 6.5, 7, and 7.5) supplemented with 10 µM valinomycin and 10 µM nigericin for 5 min. Alternatively, the cells were incubated for 30 min at 37°C in the medium containing an 1:1.000 dilution of pHrodo™ Green AM Intracellular pH Indicator (Thermo Fisher Scientific, P-35373) and an 1:100 dilution of PowerLoad™ concentrate. The cells were subsequwently washed with Live Cell Imaging Solution and analyzed on an ImageXpress high-content platform.

### Whole-cell electrophysiology

HEK293 cells were seeded on 12 mm^2^ glass coverslips in standard 24-well tissue culture plates and were allowed to grow for 24-48 h before transfection. Transfection involved the use of 0.6 μg of total DNA, 2 μL of Turbofect^TM^, and 100 μL of serum-free DMEM per well. Manual patch-clamp recordings of transiently transfected HEK293 cells were carried out in whole-cell configuration. The extracellular/bath solution contained 140 mM NMDG, 10 mM HEPES, 10 mM MES, 5 mM EGTA, 5 mM glucose, and 10 nM free [Ca^2+^] CaCl_2_ with or without 1 mM MgCl_2_. The solution was adjusted to either pH 4.6 or 7.5 with MSA. The intracellular/pipette solution contained 140 mM NMDG, 10 mM HEPES, 5 mM EGTA with or without 1 mM MgCl_2_, and variable concentrations of CaCl_2_ to reach either 10 nM or 1 μM free [Ca^2+^]. The pH was adjusted to 7.5 with MSA. The total calcium concentration at each pH was calculated to maintain the indicated amount of free calcium using https://somapp.ucdmc.ucdavis.edu/pharmacology/bers/maxchelator/CaMgATPEGTA-TS.htm. Recording glass pipettes, with resistances in the range of 5-10 MΩ were pulled and polished. Electrophysiological recordings were performed using an EPC10 patch-clamp amplifier (HEKA, Lambrecht, Germany), controlled by PatchMaster software (HEKA Elektronik). The compensation circuit of the EPC-10 amplifier cancelled fast and slow capacitive transients. In all the experiments, a step protocol was used: 400 ms voltage steps from −100 to +60 mV in 20 mV increments were applied every 5 s for a total of 13 steps, and the holding potential was 0 mV. Digitized and filtered (40 kHz and low-pass filter frequency of 2.9 kHz) current amplitudes at the end (last 5%) of each step, recording from −100 to +60 mV were extracted and normalized to the cell size. All the experiments were conducted at room temperature (22-23°C) and the data were analyzed with GraphPad Prism 10.0.2.

### Cell death

Cells (5000 cells/well) were plated on 96-well plates, and 24 h later, they were treated as indicated. After th eindicated times, the cells were incubated with 0.2 µg/ml propidium iodide or 170 nM SYTOX™ Green Nucleic Acid Stain and 2.5 µg/ml Hoechst-33342 for 10 min at 37℃, and cell death was analyzed using a Celigo Imaging Cytometer (Nexcelom Bioscience) according to the manufacturer’s instructions.

### Preparation of FeDEX solution

FeCl_2_ (0.38 g) and FeCl_3_ (0.73 g) were dissolved in 2.5 ml H_2_O each. After the two obtained solutions were mixed, 2.5 ml of 30% (v/v) NH_4_OH and 25 ml of 5% (v/v) NH_4_OH were added sequentially while a magnetic stirrer was used to stir the solution for 30 mins. After the iron particles had settled, the supernatant was discarded and the iron particles were washed twice with 25 ml H_2_O and resuspended in 20 ml 0.3 M HCl with a magnetic stirrer for 30 min. Finally, we added 1 g of Dextran 40, stirred the mor for another 30 min and dialyzed the iron dextran solution with H_2_O for 24 hours at 4°C.

### FeDEX purification

Cells grown on 15 cm petri dishes were incubated in 20 ml medium containing 100 µl of FeDEX solution. The next day, we changed the medium to normal medium. After 2-3 hours of chase, the cells were harvested in PBS and washed three times in SuMa buffer (10 mM HEPES; 0.21 M mannitol; 0.07 M sucrose; pH 7.5). The cells were then resuspended in SuMa4 buffer (SuMa 2, 1 µl Benzonase, protease inhibitor) at a concentration of 30×10^6^ cells/ml and homogenized by a gentleMACS Dissociator machine or syringe with a 25G needle. The resulting samples were pelleted by 2×10 min centrifugation at 1500 g. The supernatant was collected as the light membrane fraction (LMF). To further enrich lysosomes containing iron dextran particle, the LMF were passed through MS columns (Miltenyi Biotec, 130-042-201) and the columns were washed three times (once with SuMa4 and twice with SuMa2 (SuMa, 0.5 mM DTT, 0.5% fatty acid-free BSA in SuMa buffer)). Finally, the lysosomal fractions were eluted with 600 μl of SuMa2 from the columns, which were detached from the magnetic stand.

### Statistical analysis

Unless otherwise indicated, the statistical analyses were performed using GraphPad Prism 9 or GraphPad Prism 10 software. Data are shown as means ± SD unless stated otherwise. The specific statistical tests applied are given in the respective figure legends, where statistical significance is given by *P < 0.05, **P < 0.01, and ***P < 0.001; n.s., not significant.

## Acknowledgments

We thank Shawn M. Ferguson for providing valuable reagents, and Dianna Skousborg Larsen, Louise Vanderfox and Tiina Naumanen Dietrich for the technical assistance.

## Funding

This work was supported by the Danish National Research Foundation grant DNRF125 to M.J., the Danish Cancer Society grants R269-A15695 and R352-A20587 to M.J., the Novo Nordisk Foundation grant NNF19OC0054296 to M.J, and DFG GR4315/2-2, DFG GR4315/7-1, SFB/TRR152 P04, and SFB1328 A21 to C.G.

## Author contributions

Conceptualization: BL, MJ

Methodology: BL, KM, MJ

Investigation: RC, BL, DJ, LK, VK, BW

Visualization: RC, BL, DJ, LK, VK, BW, YK, AP

Supervision: BL, KM, CG, MJ

Writing—original draft: BL, MJ

Writing—review & editing: RC, BL, DJ, LK, VK, KM, CG, MJ

## Competing interests

The authors declare that they have no competing interests.

## Data and materials availability

All the data needed to evaluate the conclusions in the paper are present in the paper and the Supplementary Materials. Cell lines, plasmids and data sets generated during the current study will be available from the corresponding authors upon reasonable request.

## Notes

### Competing Interest Statement

The authors have declared no competing interest.

### Summary of Updates

Lines 365-368 have been revised to properly interpret data in figure 6 and to cite a recent manuscript that identified TMEM165 as a lysosomal CA2+ importer.

