## Extended Data Figures 1-5 for "TMEM165 replenishes lysosomal Ca^2+^ stores, protects cells against Ca^2+^ overload, and mediates Ca^2+^-induced lysosomal H^+^ leakage"

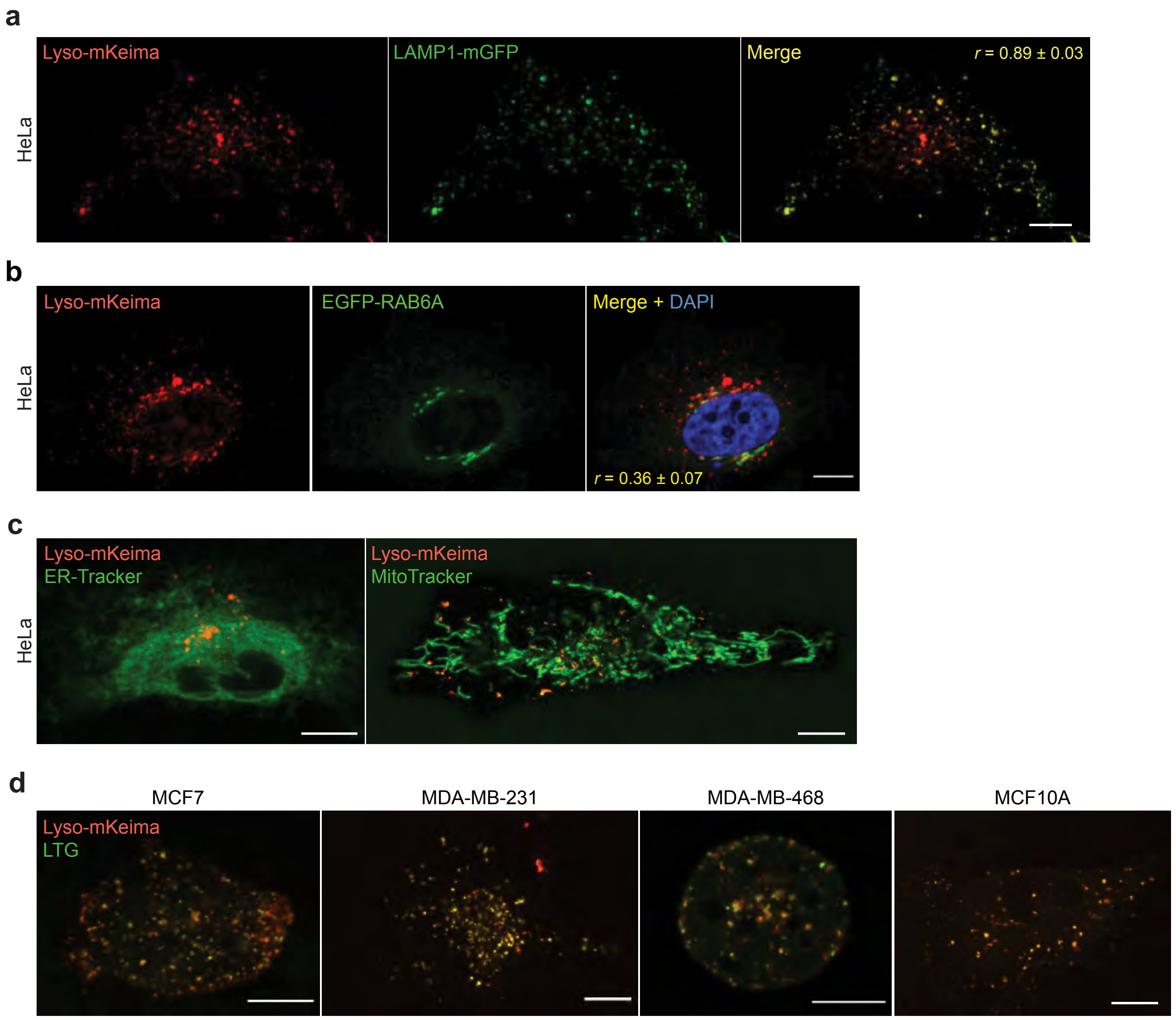


**Extended Data Figure 1. Lyso-mKeima co-localizes with lysosomal markers**

a. Representative (n = 3) confocal images of HeLa-Lyso-mKeima cells. r, Pearson's colocalization coefficient (n =20). Scale bars, 10 µm.
b. Representative (n = 3) confocal images of live HeLa-TMEM165-mCherry cells transfected with TGN markers. DNA is visualized by Hoechst staining. r, Pearson's colocalization coefficient (n=20). Scale bar, 10 µm.
c. Representative (n = 3) images of live HeLa-Lyso-mKeima cells stained with ER-tracker Green (left) or Mito-Tracker Green (right). Scale bar, 10 µm.
d. Representative (n = 3) images of Lyso-mKeima MCF7, MDA-MB-231, MDA-MB-468 or MCF-10A cells stained with Lysotracker Green. Scale bar, 10 µm.

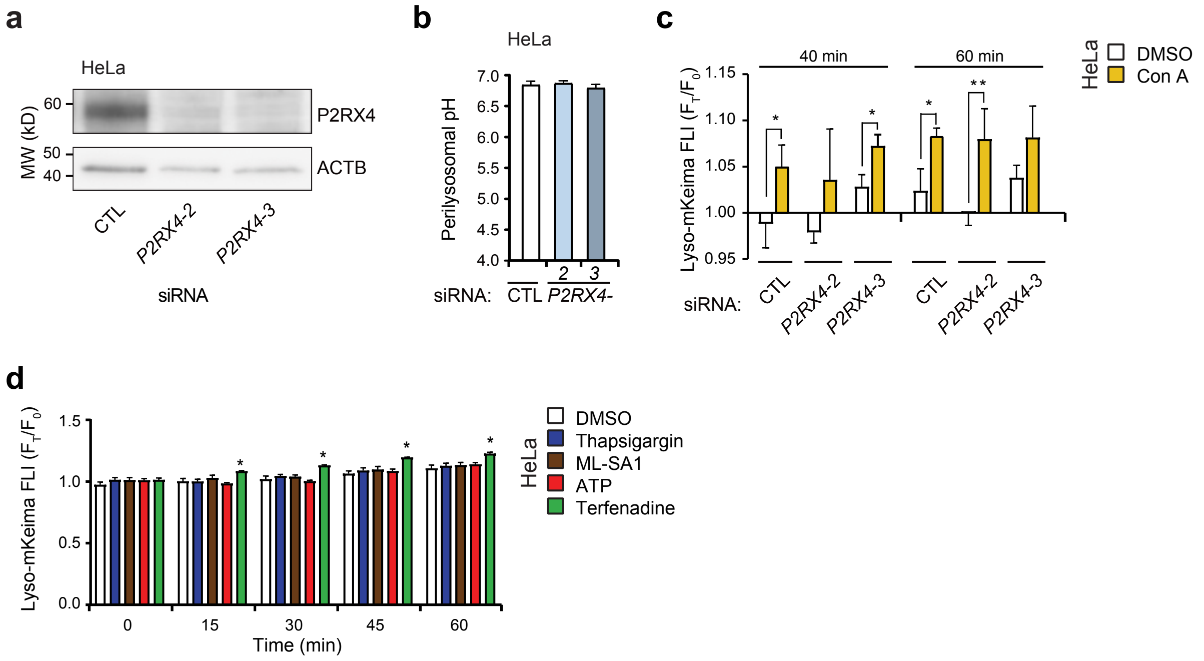


**Extended Data Figure 2. Constitutive lysosomal H+ leak is P2RX4-independent**

a. Representative (n=3) immunoblots of indicated proteins in HeLa cells treated with indicated siRNAs for 72 h.

b. Perilysosomal pH analyzed by Lyso-mKeima FLI in HeLa cells treated with indicated siRNAs for 72 h.

c. mKeima FLI in Hela-Lyso-mKeima cells treated for 72 h with CTL or P2RX4 siRNAs and for the last 40, 60 min with either DMSO or 10 nM Con A.

d. mKeima FLI in Hela-Lyso-mKeima cells treated with DMSO, thapsigargin, ML-SA1, ATP or terfenadine for the last 15, 30, 45 or 60 min.

Error bars, SD of three independent experiments with ≥10 randomly chosen cells analyzed in each sample. *, $P < 0.05$; **, $P < 0.01$; ***, $P < 0.001$ as analyzed by one-way Anova (b) or two-way Anova (c, d) with Tukey (b) or Dunnett (c, d) multiple comparison.

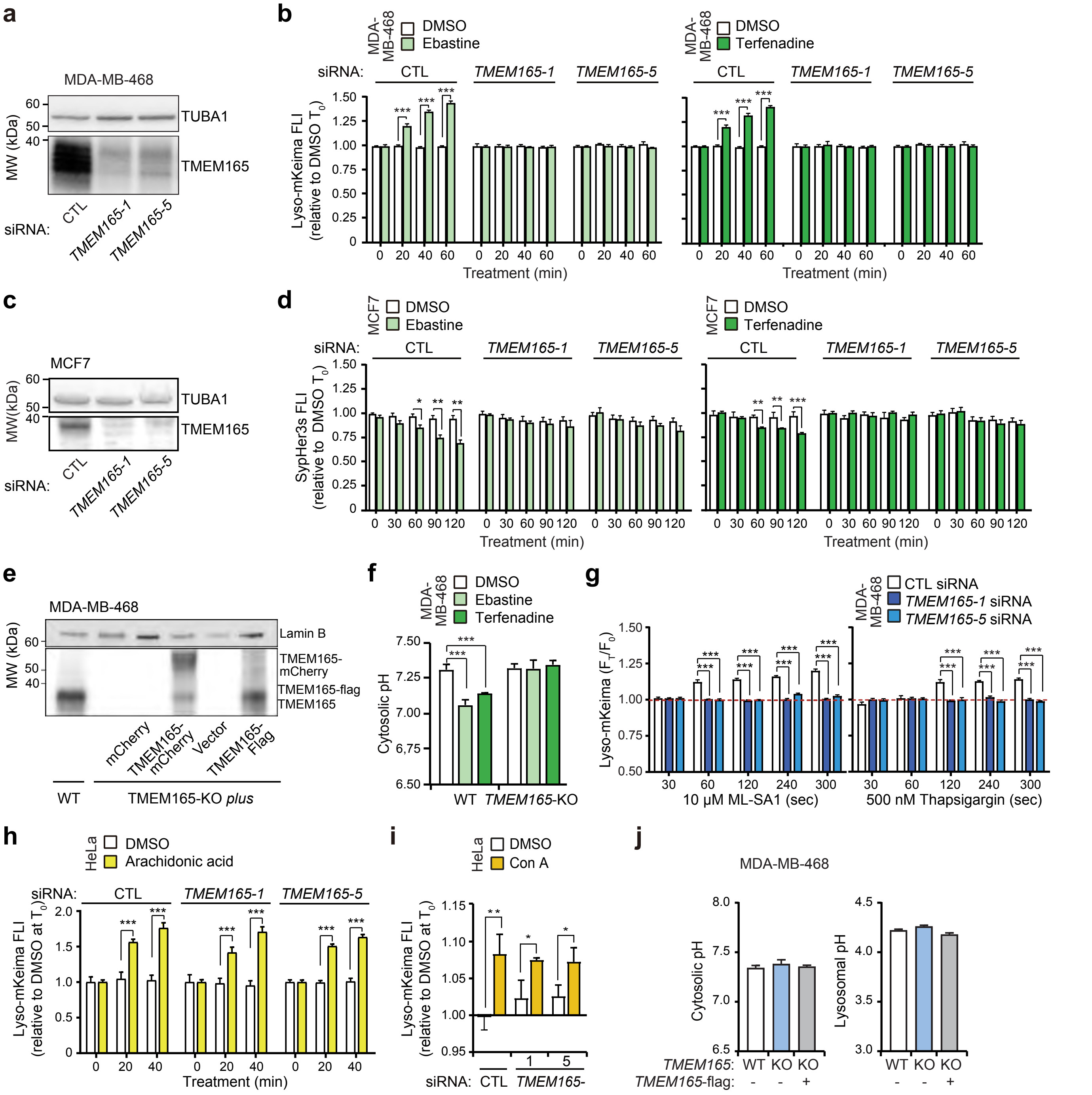


**Extended Data Figure 3. Ca2+-induced lysosomal H+ leak is mediated by TMEM165**

a. Representative (n=3) immunoblots of indicated proteins in MDA-MB-468 cells treated with indicated siRNAs for 72 h.

b. mKeima FLI in MDA-MB-468-Lyso-mKeima cells treated for 72 h with indicated siRNAs and with 15 µM ebastine (left) or 6 µM terfenadine (right) for the last 20, 40 or 60 min.

c. Representative (n=3) immunoblots of indicated proteins in MCF7 cells treated with indicated siRNAs for 72 h.

d. SypHer3s FLI in MCF7 cells treated for 72 h with indicated siRNAs and with 15 µM ebastine (left) or 6 µM terfenadine (right) for the last 30, 60, 90, 120 min.

e. Representative (n = 3) immunoblots of TMEM165 in indicated MDA-MB-468 clones.

f. Cytosolic pH analyzed by pHrodo-AM FLI in WT and TMEM165- KO MDA-MB-468 clones treated with DMSO, 6 µM terfenadine, or 15 µM ebastine for the last 1 h.

g. mKeima FLI in MDA-MB-468-Lyso-mKeima cells treated for 72 h with indicated siRNAs and with ML-SA1 (left) or thapsigargin (right) for the last 0- 300 sec.

h. mKeima FLI in HeLa-Lyso-mKeima cells treated for 72 h with indicated siRNAs and with Arachidonic acid for the last 20 or 40 min.

i. mKeima FLI in Hela-Lyso-mKeima cells treated for 72 h with CTL or TMEM165 siRNAs and for the last 60 min with either DMSO or 10 nM Con A.

j. Cytosolic pH analyzed by pHrodo-AM FLI (left) and lysosomal pH analyzed by LysoSensorTM Yellow/Blue FLI (right) in indicated MDA-MB-468 clones.

Error bars, SD of three independent experiments with ≥10 randomly chosen cells analyzed in each sample. *, $P < 0.05$; **, $P < 0.01$; ***, $P < 0.001$ as analyzed by one-way Anova (j) or two-way Anova (b, d, f, g, h, i) with Tukey (j) or Dunnett (b, d, f, g, h, i) multiple comparison.

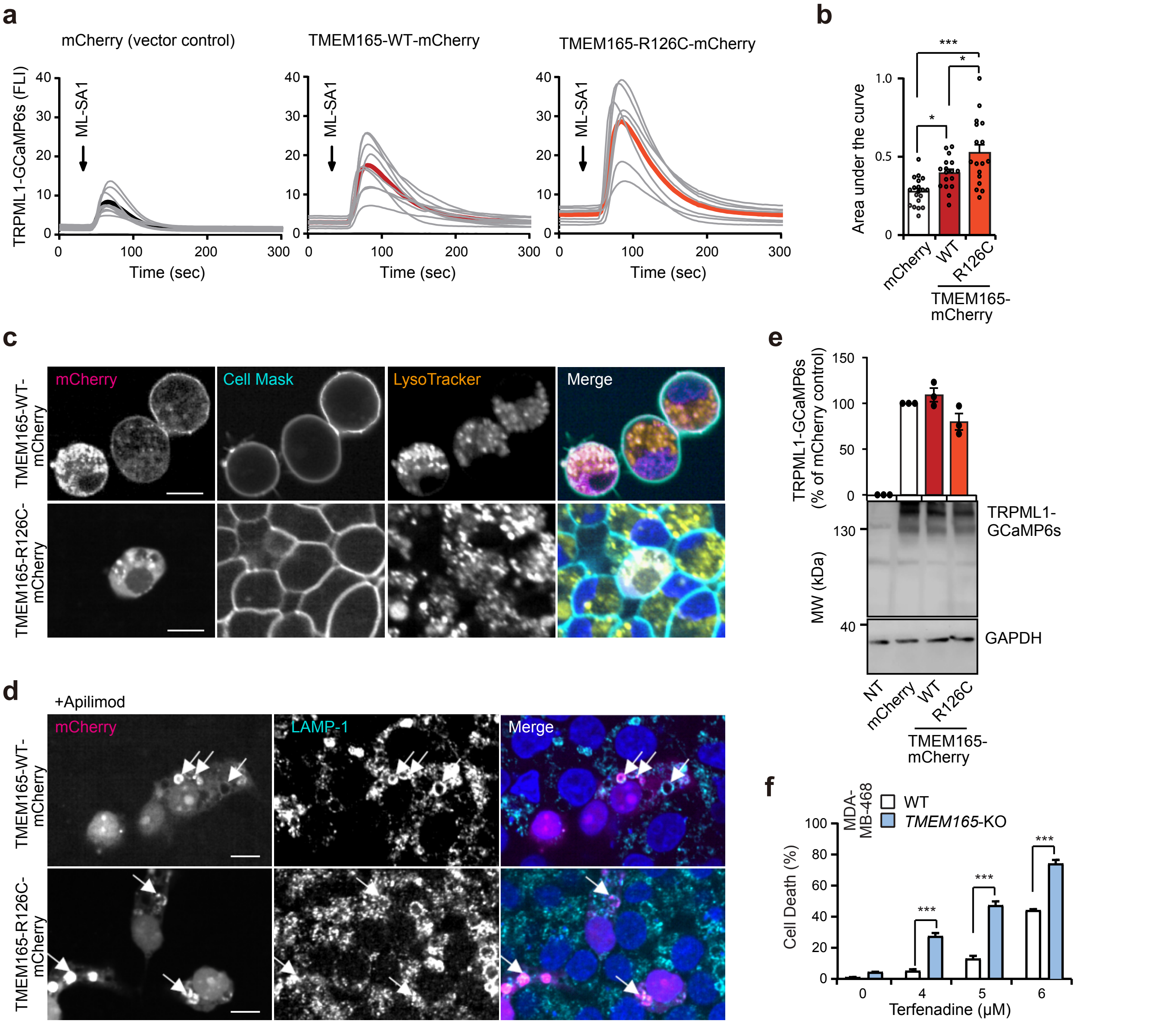


**Extended Data Figure 4. TMEM165 preserves lysosomal Ca2+ stores and protects against cytosolic Ca2+ overload**

a , b. Representative Ca2+ signals in HEK293 cells transiently transfected with plasmids encoding for TRPML1-GCaMP6s and either mCherry, TMEM165-WT-mCherry or TMEM165-R126C-mCherry (a). Thin and bold lines represent Ca2+ signal curves in response to 10 µM ML-SA1 in individual cells and the mean values of all cells analyzed, respectively (a). The bar chart represents mean values ± SEM the area under the curve corresponding to relative fluorescence unit (RFU) * seconds. N ≥ 17. (b).

c. Representative live images of HEK293 cells transfected as in a and stained with CellMaskTM green plasma membrane stain and LysoTrackerTM Deep Red and Hoechst-33342. Scale bars, 10 µm.

d. Representative confocal images of HEK293 cells transfected as in a, treated with 1 µM apilimod for 16 h, fixed, stained with LAMP1 antibodies and Hoechst-33342 and imaged at the Opera Phenix High Content Imaging System. Scale bars, 10 µm

e. Representative (n=3) immunoblots of TRPML1-GCaMP6s (anti-GFP) and GAPDH (bottom, loading control) and quantification of relative TRPML1-GCaMP6s protein levels (top) in HEK293 cells transfected as in a. Error bars, SD of three independent experiments.

f. Death of indicated MDA-MB-468 cell clones treated with terfenadine for 24 h. Cells were stained with propidium iodide (dead cells) and Hoechst-33342 (total cells) and cell death was analyzed by Celigo Imaging Cytometer. Error bars, SD of three independent experiments with ≥ 10000 randomly chosen cells analyzed in each sample.

*, P < 0.05; **, P < 0.01; ***, P < 0.001 as analyzed by one-way ANOVA with Tukey's multiple comparisons (b) or two-way Anova with Dunnett's multiple comparison (c).

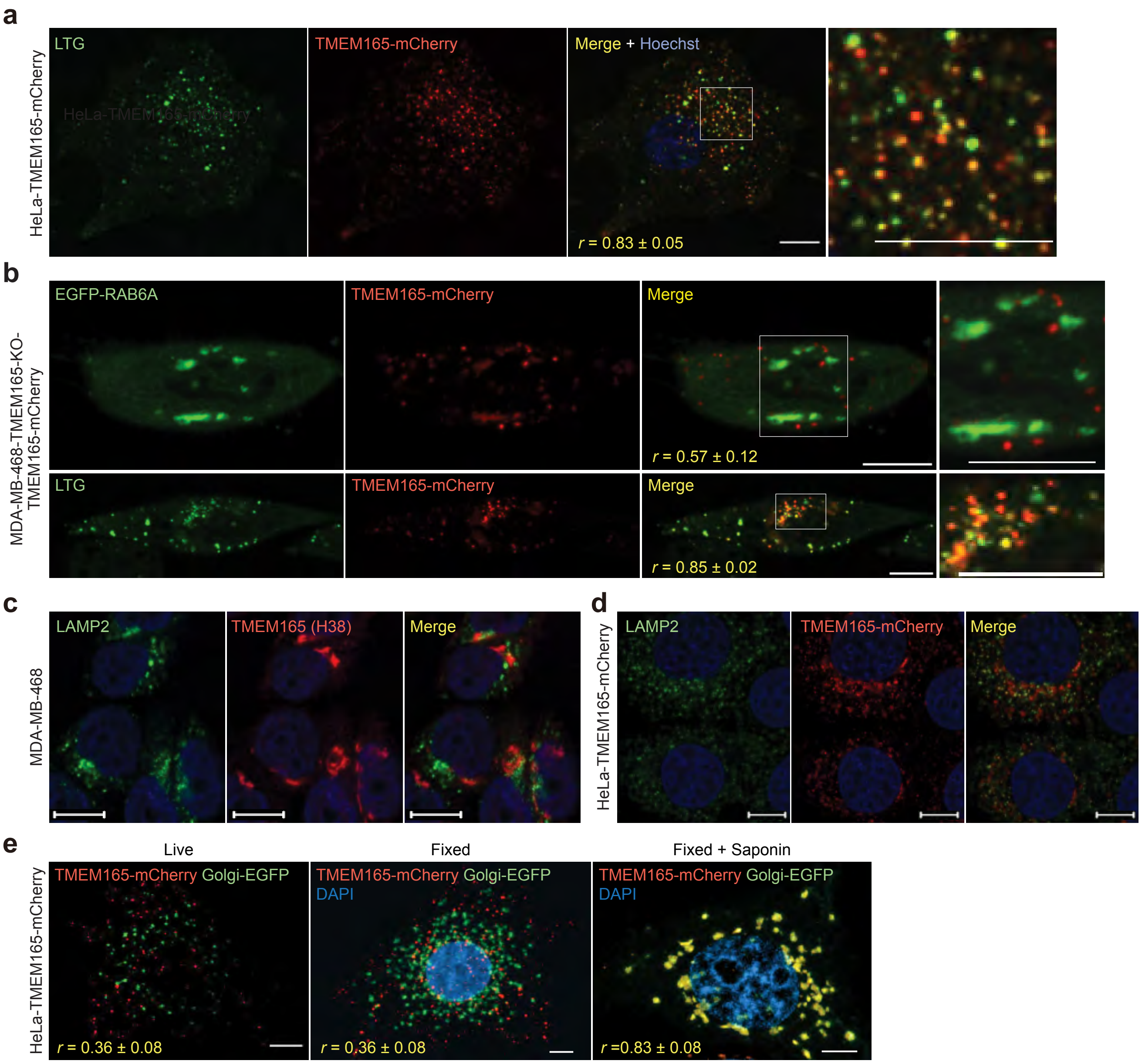


**Extended Data Figure 5. TMEM165 localizes to lysosomes and Golgi**

a. Representative (n > 3) confocal images of live HeLa-TMEM165-mCherry cells stained with Lysotracker® Green. DNA is visualized by Hoechst staining. White squares mark the area shown in enlarged images (right). r, Pearson's colocalization coefficient (n=20). Scale bar, 10 µm.

b. Representative (n = 3) confocal images of live MDA-MB-468 TMEM165-KO-TMEM165-mCherry cells transfected with TGN markers (top) or stained with Lysotracker® Green (bottom). White squares mark the area shown in enlarged images (right). r, Pearson's colocalization coefficient (n ≥ 10). Scale bars, 10 µM.

c. Representative (n = 3) confocal images of MDA-MB-468. Scale bars, 10 µM.

d. Representative (n = 3) confocal images of HeLa-TMEM165-mCherry cells. Scale bars, 10 µM.

e. Representative (n = 3) 3D confocal images of live (left) and fixed (middle and right) HeLa-TMEM165-mCherry cells transfected with Golgi-EGFP with or without permeabilization with saponin. r, Pearson's colocalization coefficient (n ≥ 10). Scale bars, 5 µM.
